# DeepFRET: Rapid and automated single molecule FRET data classification using deep learning

**DOI:** 10.1101/2020.06.26.173260

**Authors:** Johannes Thomsen, Magnus B. Sletfjerding, Stefano Stella, Bijoya Paul, Simon Bo Jensen, Mette G. Malle, Guillermo Montoya, Troels C. Petersen, Nikos S. Hatzakis

## Abstract

Single molecule Förster Resonance energy transfer (smFRET) is a mature and adaptable method for studying the structure of biomolecules and integrating their dynamics into structural biology. The development of high throughput methodologies and the growth of commercial instrumentation have outpaced the development of rapid, standardized, and fully automated methodologies to objectively analyze the wealth of produced data. Here we present DeepFRET, an automated standalone solution based on deep learning, where the only crucial human intervention in transiting from raw microscope images to histogram of biomolecule behavior, is a user-adjustable quality threshold. Integrating all standard features of smFRET analysis, DeepFRET will consequently output common kinetic information metrics for biomolecules. We validated the utility of DeepFRET by performing quantitative analysis on simulated, ground truth, data and real smFRET data. The accuracy of classification by DeepFRET outperformed human operators and current commonly used hard threshold and reached >95% precision accuracy only requiring a fraction of the time (<1% as compared to human operators) on ground truth data. Its flawless and rapid operation on real data demonstrates its wide applicability. This level of classification was achieved without any preprocessing or parameter setting by human operators, demonstrating DeepFRET’s capacity to objectively quantify biomolecular dynamics. The provided a standalone executable based on open source code capitalises on the widespread adaptation of machine learning and may contribute to the effort of benchmarking smFRET for structural biology insights.

## Introduction

Single molecule Förster resonance energy transfer (smFRET) combined with TIRFm (total internal reflection fluorescence microscopy) is a key powerful method to study the structure of biomolecules and provide a dynamic perspective in structural biology (1). Capturing the real-time readouts of nanometer scale distances of individual biomolecules by smFRET allows the direct observations of dynamics, interactions and intermediates of stochastic non accumulating events, as well as dynamic equilibria between unsynchronized molecules, all of which are obscured in ensemble averaging techniques (2–10). The high fidelity and proficiency, of smFRET established it as a key toolbox for the accurate characterization of mechanisms, biomolecular interactions function and even structures of biomolecules (11–15), under both *in vitro* (16–18) and *in vivo (19, 20)* conditions. Despite its great quantitative utility and profound impact for structural biology, smFRET is not a direct imaging modality and data treatment for extracting quantitative dynamic information rely on multiple layers of preprocessing: raw image treatment, trace selection and data analysis. Raw image treatment (4, 5, 10, 21, 22) and data analysis of the selected smFRET traces is in general well-standardized and relies on well-defined methodologies with strong theoretical backing (21, 23).

The actual trace selection is time consuming, but crucial due to the presence of undesired phenomena at the single molecule scale, such as sample aggregation, fluorescent contaminants, incomplete or incorrect sample labeling, complex photophysical behaviors and high noise, to mention a few (3, 8, 24). Existing software (3–5, 10, 21, 22) by single molecule labs can simplify the tedious and time consuming selection of traces, and were recently expanded to large data sets (4), albeit requiring some form of manual supervision and hyper-parameter tuning by an expert user. This need for human intervention could potentially be subjected to cognitive biases especially by less experienced users and could limit the expansion of smFRET to classic biology labs. The increasing expansion of smFRET to structural biology labs would benefit from rapid and benchmarked methodologies, reproducible across laboratories, with minimal human intervention. This is highlighted by several initiatives to standardise the smFRET field (1, 3, 22, 25, 26).

Recent advances in machine learning (ML) and specifically deep learning (DL) (27), have radically improved our capacity to access and extract information from abstract and noisy inputs independently of human interventions as we (28) and others have shown (29–36). DL implementations are providing high level robust performances and have been successfully used to analyze and augment a wide range of the fluorescence microscopy analysis pipeline including assessing microscope image quality (37), in-silico cell labeling (31), single cell morphology analysis (32, 34), detecting single molecules (38) and linking smFRET experiments with molecular dynamics simulations (39), amongst others (29–36).

Deep learning-based analysis has several advantages over other approaches: It recognizes abstract patterns and learn useful features directly from the raw input data which allows implementation of analysis routines that don’t require extensive data preprocessing or empirically defined rules and thus offer reproducible and less opinionated evaluation of single molecule data; It is significant faster than human annotation for large single molecule data sets; it comes close to, or outperforms human performance; and its performance is increased when increasing data set size constituting an ideal case for evaluating the large data sets obtained from single molecule data (29–36). Especially important are convolutional DNN which learn how to best recognize particular aspects of the given data through several rounds of optimization. The network then classifies data into predefined classes based on the provided training labels. While the training of a DNN is generally a computationally intensive process, once trained the final model can easily be shared and used for making predictions at almost no computational cost to end users.

Here we provide DeepFRET, an all-inclusive analysis software with a pre-trained DNN at its core, for rapid, objective and accurate assessment of smFRET data for quantifying biomolecular dynamics. The fully automated analysis software operates with minimal crucial human intervention and requires only a threshold on the data quality confidence, as an initial step, so as to output detailed quantification of structural dynamic from raw images. This is attained by an intuitive and user-friendly interface that integrates and automates common smFRET analysis procedures (3–5, 10, 21, 22) from raw image analysis and background-corrected intensity trace extraction (40), to sophisticated trace classification, statistical analysis of single molecule data and production of publication quality figures of dynamic structural biology insights (see Materials and Methods). DeepFRET comes as a free-to-use standalone executable allowing end users with limited programming skills to easily operate it. A script-based version implemented entirely in Python enables experts to adjust features pipelining the analysis specified for their needs. We anticipate DeepFRET to take full advantage of the widespread digitization and open repository of smFRET data and form a reference point setting a bar for the data quality and data classification performance metrics, offering additional benchmarking the field for dynamic structural biology.

## Results

### DeepFRET software package

DeepFRET is an open source software package that implements a neural network model architecture for data evaluation integrating in a user-friendly platform all common procedures for smFRET analysis (Fig. 1). The neural network model architecture used here (Fig. S1 and S2) is based on a deep convolutional neural network to recognize particular spatial features present in the data. The model first passes the data through several layers of convolutions of different lengths, and in the process learns to recognize which particular elements of a sample are present at different length scales, to best classify it correctly. This has previously been used to label time series data such as electrocardiographs (41) or electrical readouts in DNA sequencing (42). Additionally, we added a long short-term memory (LSTM) layer after the convolutional layers, as this will also help the model to learn temporality in the data and propagate to the later frames the learned information (See Methods) (43, 44). A detailed description of the model hyperparameters and architecture can be found in the Methods section.

**Fig. 1.**
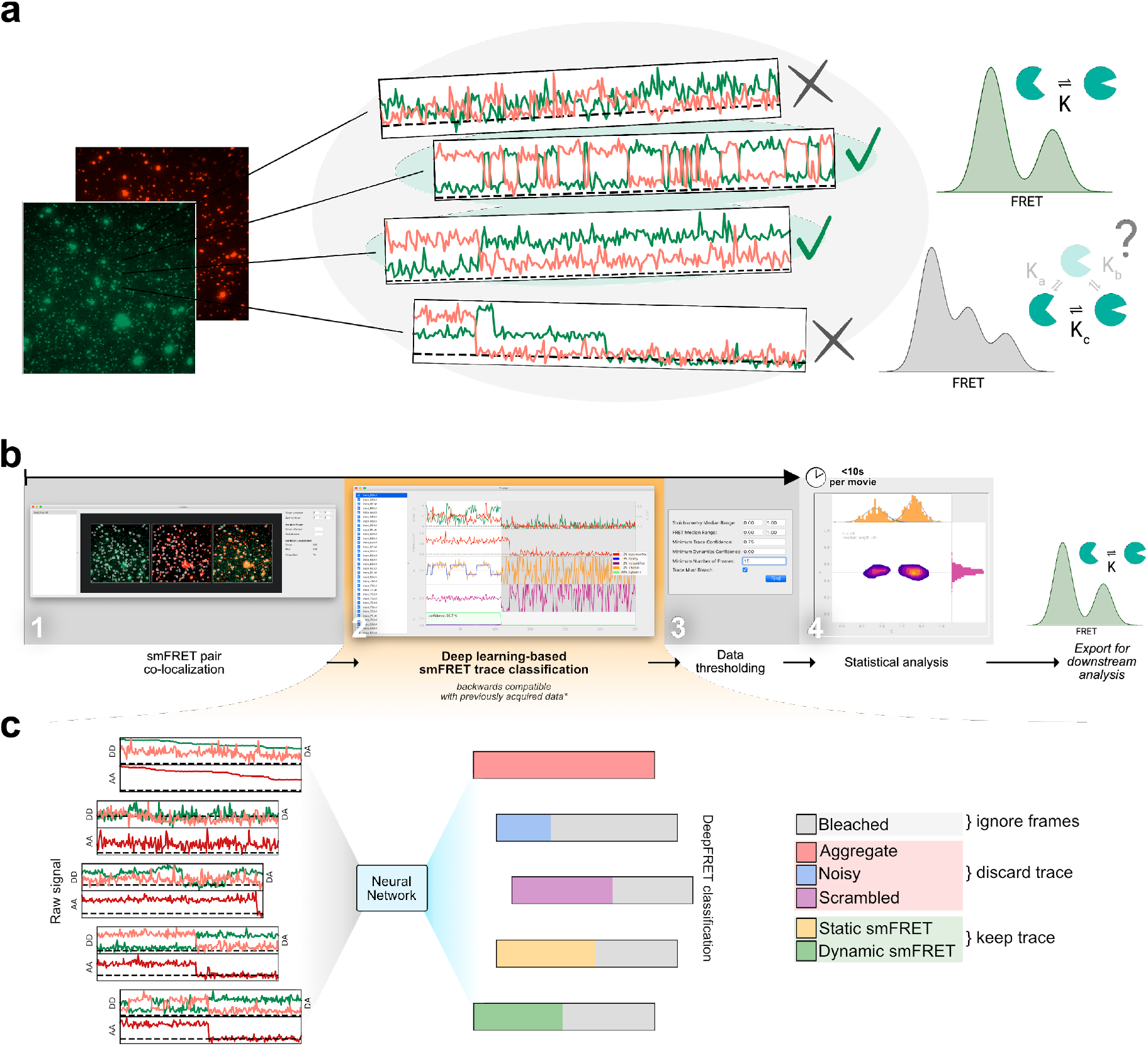
Overview of single molecule FRET evaluation and analysis using DeepFRET. **a)** Cartoon of the typical heterogenous data acquired in smFRET experiments. The limited standardization in the field for the criteria for data selection for downstream analysis may yield different structural and kinetic information. **b)** Screenshots of the provided standalone software that integrates deep learning and reduces the selection to a single user-adjustable criteria: the confidence threshold. The simple and intuitive GUI integrates all the features of our approach for rapid traces extraction from raw images to filtering of traces based on the predicted classification, treatment of smFRET data to extraction of publication quality figures **c)** End-to-end sequence classification of smFRET traces by deep learning. Raw signals of donor-donor, donor-acceptor and acceptor-acceptor intensities in the form of ASCII files can also be loaded with the DeepFRET software. The pre-trained DNN will classify individual frames to one of six different categories: bleached, static smFRET, dynamic smFRET, aggregate, noisy and scrambled. A final smFRET confidence score is calculated, depending on each of the categories, that is used for threshold.

To ensure that the predictions of DeepFRET would generalize to a wide range of experimentally observable behaviors independently of biological systems or experimental conditions, we provide a fully pre-trained DNN model. The implemented DNN is pre-trained on 150.000 simulated traces that uniformly sample all possible FRET states, their respective lifetimes and occupancies, as well as all possible noise levels, ensuring that the data represents all theoretically possible configurations (See Fig. S3-S5 Materials and Methods for software and algorithms). As such DeepFRET does not require the selection of any direct initial guesses of FRET values or user defined parameter pretraining. We do however provide both a script-based method for simulating smFRET data, as well as a simple graphical interface for expert end-users to adjust simulation distribution parameters (see See Fig. S6 and Materials and Methods) if needed (e.g. for specific circumstances or stricter criteria). This offers experts the possibility to benchmark the impact of e.g. one’s own sorting criteria, noise and optical correction factors.

We built DeepFRET to treat both Alternative Laser Excitation (ALEX) and non ALEX FRET data. DeepFRET imports raw microscope images and performs colocalization of the two channels, to extract background corrected intensity traces of DD (donor excitation; donor emission), DA (donor excitation; acceptor emission), their respective stoichiometry, and in the case of ALEX data, also AA (acceptor excitation; acceptor emission) (see Fig. 1a, and Fig. S4). Alternatively, one can directly load and process previously-obtained time-traces without their associated videos.

For a given time trace the DNN predicts and outputs six softmaxed probabilities *p_i_* to each time frame (Fig. S5 and Methods), representing the six categories it has been trained to recognize: bleached (B), static smFRET within the experimental time frame (S), dynamic smFRET (D), aggregate (A), noisy (N), and all other types of non treatable data smFRET data defined here as scrambled (X) (Fig. S5 and S7). Both static and dynamic traces are included for further analysis. Given these probabilities, which sums to one, a simple sliding window then searches for frames predicted by the DNN to be bleached (*p_B_ > 0.5*, see Fig. S5, S7, S8 for evaluation accuracy, and blinking exclusion). When bleaching is found the rest of the trace is removed so as to exclude the photobleached frames part of a trace from further analysis. If the trace still contains a minimum number of viable frames (here set to 15, but adjustable), the probabilities are summed up over all remaining time frames for each of the five remaining categories and divided by the number of frames for normalisation (see Materials and Methods and Fig. S5, S7, S8). We define the summaries of the combined static and dynamic trace scores as the “DeepFRET score”, representing the DNN model confidence that a trace is truly smFRET. The user-friendly interface displays all the categories and their associated probabilities, and offers the option for expert users to manually revise the classified traces.

If the DeepFRET score is above the user defined threshold, the trace is accepted for subsequent analysis (see Fig. 1b, Fig. 1c and Materials and Methods). Subsequent analysis involves two-channel fitting of idealized FRET traces using Hidden Markov modelling *HMM* (using the open-source package pomegranate); data and statistical evaluation of the abundance of FRET states and lifetimes; application of correction factors; and transition density plots (see Fig. S6, S9, S10, S11). The number of underlying FRET distributions is automatically determined using Bayesian information criterion (BIC), offering the unbiased analysis of distribution of biomolecular distances (See Fig. S6, S9-S11). All data can be directly exported in publication quality figures or extracted as data for user specific analysis if required.

### Performance of DeepFRET

To test our DeepFRET performance in practice we initially compared it with commonly used threshold values. We simulated 200 ground-truth smFRET traces and merged them with a dataset containing 5000 random, non-smFRET traces (too noisy, aggregates of multiple molecules, aberrant single molecule behavior. see Methods for parameter descriptions). The obtained overall FRET distribution is akin to what one would observe experimentally before any preprocessing of smFRET data on proteins (Fig. 2a). Common procedures for pre-selecting valid data for treatment often rely on an initial automatic threshold for discarding this large fraction of non-smFRET data (see Fig. 1a). This is based on any number of combinations of the anticorrelated signal of the donor and acceptor, fluorophore bleaching, noise levels, or certain ranges of fluorophore stoichiometry, if recorded using ALEX methods (3, 45, 46). We first removed photobleaching and then accepted or rejected traces based on commonly used thresholds of median stoichiometry and max intensity (but not anticorrelation, see Materials and Methods) without any manual post-inspection of the data. Fig. 2b displays ground truth distribution (green) and the distribution of the accepted traces (pink) for varying the above thresholds. We recovered a poorly-defined FRET distribution, that even at the tightest threshold does not recapitulate the underlying ground-truth two-state conformational equilibrium. We calculated the common model evaluation metrics “precision” and “recall” (see Materials and Methods) to quantify the quality of the predictions. The precision and recall, though improved by tightening the threshold, remain around 0.22 and 0.40, in the best case for simple thresholding (Fig. 2b). The fact that out of 366 selected traces only 80 were true positive, while 286 were false positive and 120 false negative, highlights the need for human intervention as many traces are indistinguishable with simple statistical characterization (selected examples shown in the examples below the histograms (Fig. 2b).

**Fig. 2.**
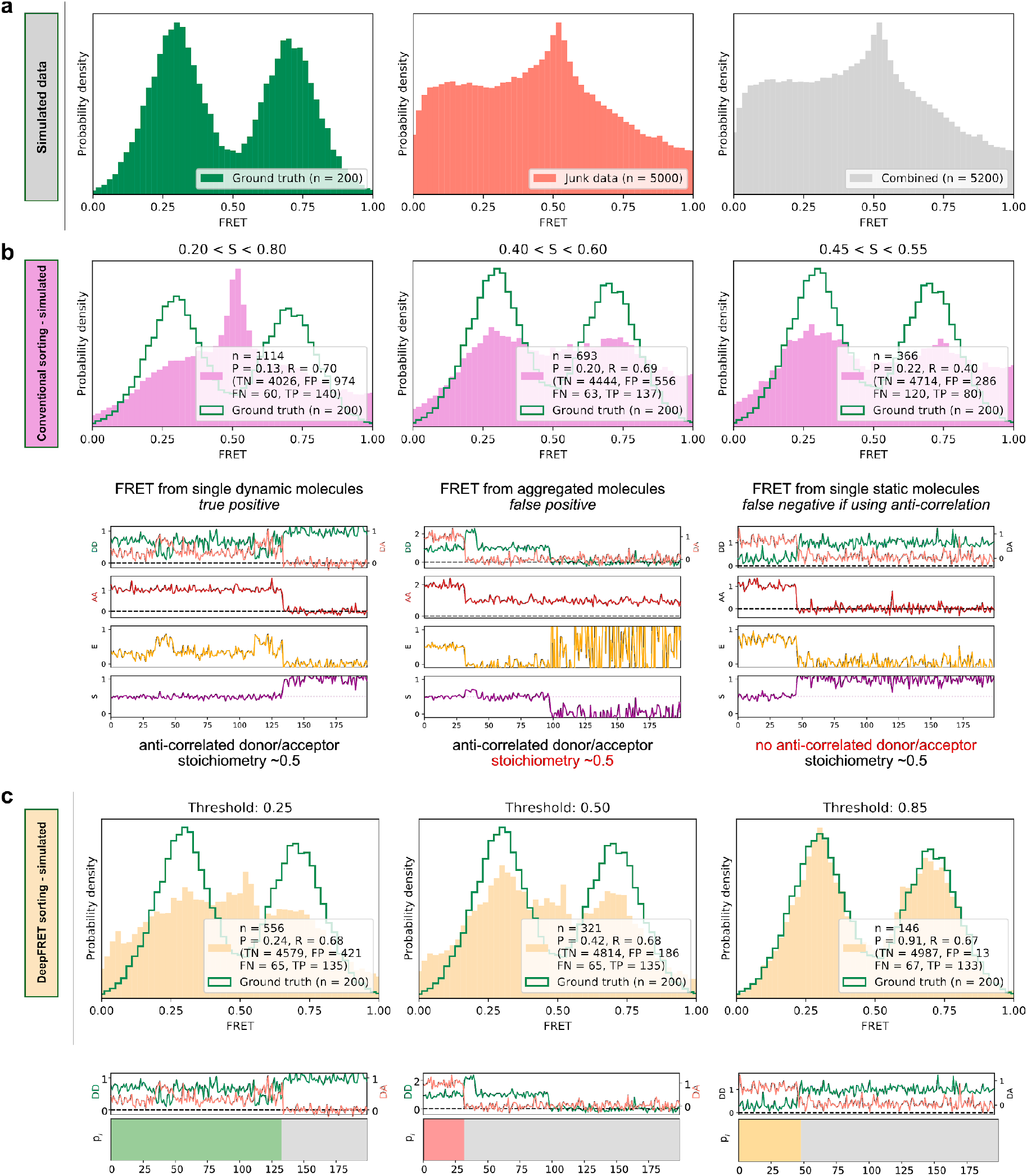
High quality FRET data evaluation. **a)** Simulated dynamic smFRET traces transitioning between FRET states 0.3 and 0.7 (left, “ground truth”) were mixed with a larger number of traces not showing smFRET (center). The overall distribution (right, “combined”) shows how the desired data can be drowned out in non-smFRET contaminant traces. The distribution would correspond to a raw distribution as extracted from raw image analysis of smFRET on proteins before any trace selection **b)** Automatic selection of data based on median stoichiometry, single molecule intensity and bleaching. The number *n* designates the number of traces accepted by the model. Tightening the selection thresholds results in slight improvement of the poor overlap of the selected data with ground truth data, highlighting the need for a time consuming and prone to potential cognitive biases human intervention. **c)** Automatic classification of all traces of the combined set by DeepFRET, based only on DeepFRET score threshold variation. Even at a low threshold DeepFRET selection follows the ground truth data. Increasing the score threshold further increases the fidelity of data selection. DeepFRET correctly assigns the dynamic, bleaching and aggregate behavior on the same smFRET traces as in (b) (see Fig. S4 for more data). The single user adjustable score threshold outperforms commonly used thresholds offering rapid, cross-lab reproducibility and fully automatic data treatment. P: precision, R: recall, TN: true negatives, FP: false positives, FN: false negatives, TP: true positives.

DeepFRET on the other hand allowed the high fidelity recovery of the underlying ground truth distribution reaching precision of 0.91 when setting a DeepFRET score of 0.85 (Fig. 2c) without the need of human intervention. The virtually identical FRET distributions, matching the ground truth data, that are derived for practically the entire spectra of score thresholds (0.25 to 0.85) show no systematic biases originating from data evaluation and illustrate the minimum impact of human interventions when using DeepFRET (see also Fig. S12). As expected the fidelity of DeepFRET pertained to correctly identifying single or complex multistate FRET distributions (see Fig.2 and Figs. S12, S13) reaching precision 0.91 as compared to just 0.22 for standard threshold setting in the absence of further human intervention. The practically identical precision and recall for single, double or triple, state FRET distributions independently of threshold further support the wide applicability to multiple biological systems.

Quantification of precision and recall of the selection for various DeepFRET score thresholds displays the trade offs in recovering high fraction of useful data. (see Fig. S12). Thresholding data with scores in the regime 0.8-0.9 appears optimal for maintaining sufficient and high fidelity data (Fig. S12). Based on these data we suggest a score threshold of 0.85 as optimal for maintaining high precision at reasonable recall values. Depending on data sets users may need to adjust the threshold. The power of DeepFRET is further highlighted by the classification for the traces that were assigned as false negative and false positive by commonly accepted thresholds (traces in Fig. 2B and Fig. 2C) (see also Fig. S4 and Fig. S14). In summary, the fidelity of classification accuracy appears to significantly supersede currently used simple thresholding, without human interventions. This was achieved in a fraction of the time required for data classification by human operators (~1 minute for 10,000 traces on a normal laptop, as compared to potentially days for manually inspected traces). This improved classification was also achieved entirely without any preprocessing or post-inspection of data, illustrating the power of DeepFRET to operate without human interventions and its potential to benchmark the reproducibility of smFRET data acquisition methods for multiple biomolecular systems across laboratories.

We quantified and displayed using confusion matrix the discordance between the ground truth data and the data selected and classified by DeepFRET (Fig. 3). In the confusion matrices displayed in Fig.3 each row represents the predicted classification of traces while each column represents the ground truth data. The high classification accuracy for the annotation of individual frames is highlighted by the clear diagonal feature. We found similar classification performance for a DNN trained on non-ALEX FRET (by a DNN with only DD and DA inputs, which we will refer to as “ALEX-disabled”) (Fig. 3 right panels) signifying the applicability of the DeepFRET approach to both ALEX and non-ALEX FRET data. The misclassification between static and dynamic smFRET traces is practically non-existent, and consists of <3% dynamic traces being misclassified as static, for both model types. This is important for accurately quantifying the abundance of static and dynamic subpopulations within the experimental time frame, which has been shown to have a clear experimental impact (12, 47–49).

**Fig. 3.**
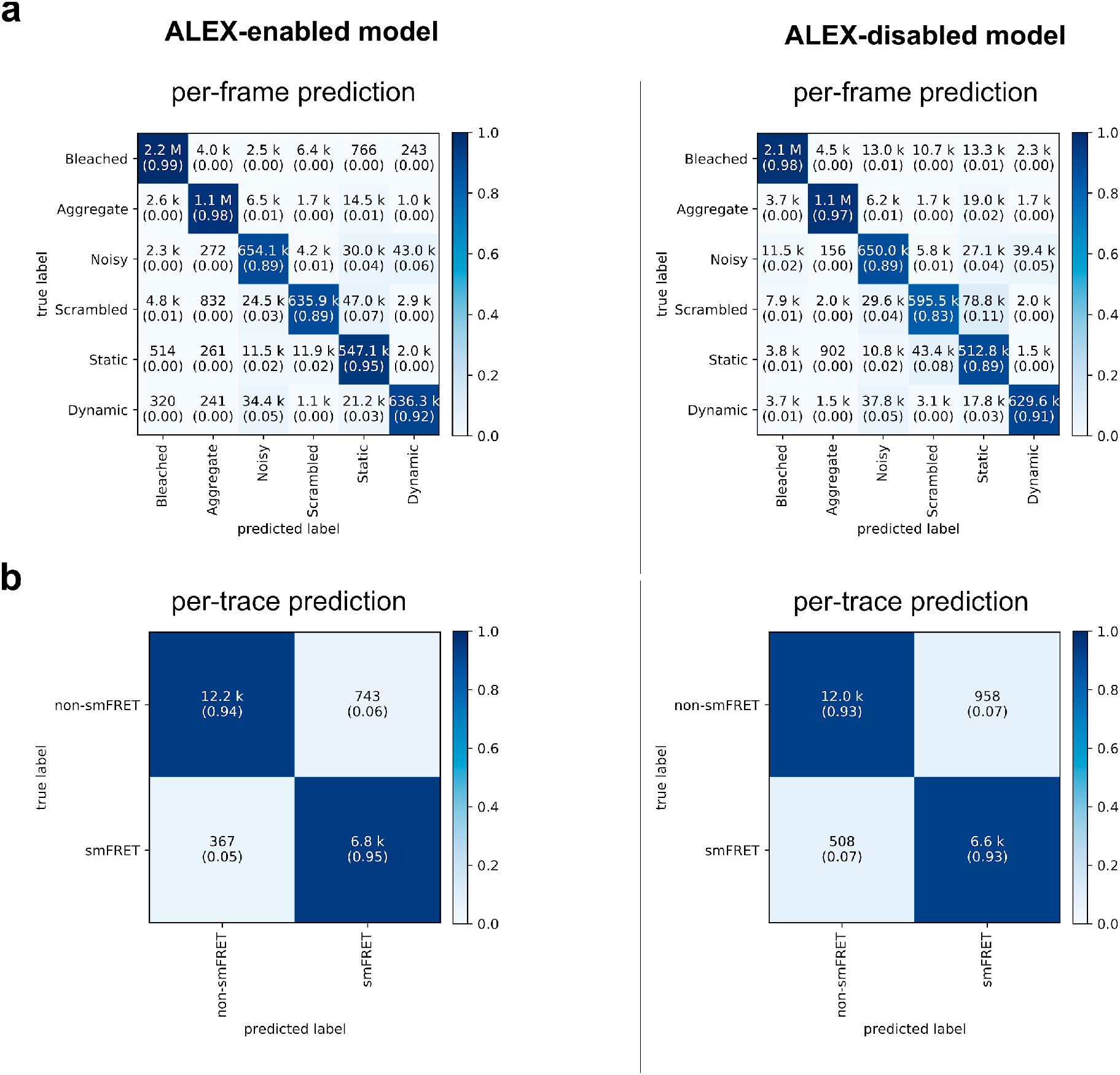
Confusion matrices of DeepFRET classification based on the ground truth data test. a) Classification accuracy of data in the 6 categories for ALEX-enabled model, or the ALEX-disabled model. The absolute number of frames is shown while the fractions for each classification is displayed in parentheses (as calculated row-wise for each true label). The diagonal percentages shows the accurate classification of DeepFRET **b)** per-trace classification accuracy, based on accepting only traces that are classified as smFRET (static/dynamic), and nonFRET data.

DeepFRET was found to classify bleached or aggregated frames with a 98% the true positives for ALEX-FRET model enabled (97% for the non-ALEX model), whereas only 89% (and 83% for ALEX-disabled) of the scrambled traces were correctly classified (see also Fig. S14 for a detailed breakdown of the precision and recall). We note that the model is trained with a noise contribution that is drawn from a normal distribution of varying width (σ between 0.01 to 0.30, multiplied by the maximum single fluorophore intensity) with a small contribution of gamma-distributed noise. As such traces with σ above 0.25 are characterised by the employed DNN as “noisy” (see Fig. S14 and Methods). In order to allow experts to accept more noisy traces, or traces with fast transition rates that may appear as noisy for a given imaging conditions, we integrated in DeepFRET a visual trace simulator. This user friendly simulator allows generation of traces with ground truth labels of traces where all parameters are tunable so as to integrate the specific needs of each lab (See Figs. S6, S11).

We found the classification accuracy of each frame to be consistent with the classification accuracy on each trace, later derived from the overall most probable class given all predictions of individual frames of a trace (Fig. 3a, Fig. 3b, Fig. S2, S6 and Methods). This is achieved by adding a bidirectional long short-term memory, LSTM, layer at the end of the DNN (Fig. S1). The LSTM layer allows coherent predictions throughout the trace and forward propagation of information detected in the first frames such as e.g fluorophore detection or bleaching, to the predictions for later frames. By collapsing the per-trace confusion matrix into a binary “smFRET” and “non-smFRET” (as shown by the cross-lines in Fig. 3b), DeepFRET was found to be very balanced overall, with a true-positive rate of 94% for smFRET traces, and a true-negative rate of non-smFRET traces (Fig. 3c), resulting in an overall balanced classification accuracy of 94% for the ALEX-enabled model and 93% for the ALEX-disabled model.

We then compared the classification accuracy of DeepFRET to the accuracy of three different human operators working with smFRET, to evaluate the feasibility of manually inspecting and making decisions about smFRET examples. We simulated 1000 ground truth traces, of which only 46 contained actual smFRET, at different, randomly chosen levels of noise. The participants were not informed about the underlying distribution, nor the true number of smFRET traces. The test revealed that the average performance of the human operators, scoring 0.76 +/− 0.10 in precision and 0.83+/−0.14 in recall, was close to the precision-recall curve of the DNN, on a relatively small dataset (Fig. S15). Notably, one participant scored slightly better than the model in both precision and recall, but spent an average of 5 s per trace, which would significantly increase data treatment time, thus making this unfeasible in a high-throughput setting. The large spread on precision and recall attained by human operators on these data furthermore suggests a large possible spread in experimental outcomes and highlights the advantages of unifying, reproducible methodologies independent of human interventions. We therefore argue that DeepFRET is equally good, or better, as careful manual inspection while offering orders of magnitude faster data evaluation.

### DeepFRET performance on real data

The model’s generalizability was subsequently demonstrated by evaluation on real experimental smFRET data previously published by us (10). The selected published dataset contains thousands of traces that included aggregates and incomplete labeled molecules, due to the low labeling efficiency. Our pre-screening (using median stoichiometry and intensity distributions) and subsequent manual inspection resulted in 214 to exhibit smFRET. Applying our trained model with a threshold of 0.85, without any other parameter tuning, recovered 228 traces, with a FRET distribution very closely matching manual selection (Fig. 4, Fig. S16). The DeepFRET score of human vs machine selection displays the importance of quantitative and reproducible assessment of trace scores (Fig. S17). The total data evaluation time of < 50 ms per trace (on a recent laptop) free of human intervention highlights the potential of DeepFRET to rapidly and reliably evaluate high throughput single molecule FRET data. Most importantly, the trace selection is deterministic, strictly relies on the score threshold and is thus independent of potential human cognitive bias. This clearly demonstrates the ability of the DNN to generalize to a completely new set of experimental data, without any prior expectations for signal-to-noise ratio, anti-correlation, underlying FRET distribution, etc. offering the possibility to rapidly analyse single molecule FRET data for structural biology insights.

**Fig. 4.**
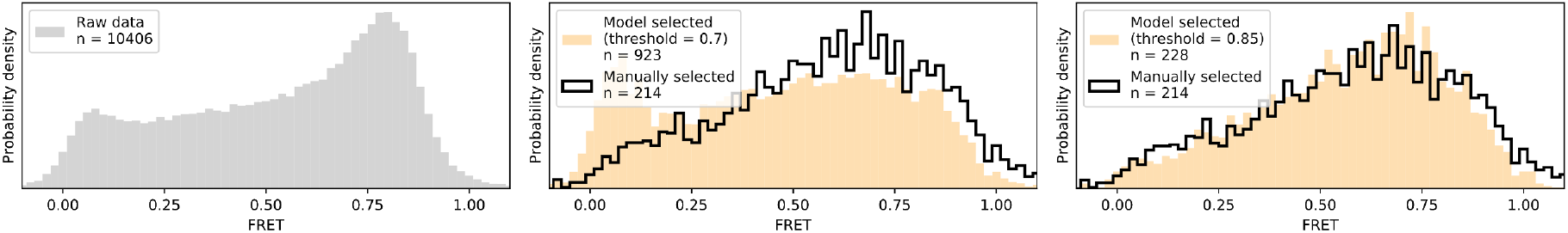
Method evaluation on real, previously published smFRET data. At DeepFRET score threshold of 0.85 a high fidelity data selection is achieved resulting in a similar distribution as compared to manual selection.

To ensure the facile operation of DeepFRET by non-machine learning experts and users without any programming skills we provided a standalone executable along with simple and detailed instructions on how to use it (see Methods). DeepFRET implements and automates in a user friendly and intuitive platform all common procedures for single molecule FRET analysis: sophisticated raw image analysis from raw.tiff files; particle and signal detection and localization; pixel intensity extraction for each individual biomolecule on both spectral channels and background corrected fluorescence and FRET trace trajectories; automatic trace classification and sorting; unbiased analysis of number of FRET states based on BIC analysis; 2-channel fitting of idealized FRET traces using HMM analysis based on calculated number of states by BIC; data and statistical evaluation of abundance of FRET states and lifetimes; application of correction factors; and transition density plots (see Figs. S9-11). DeepFRET furthermore offers interoperability and backwards-compatible trace loading from.txt files exported from the popular iSMS software package. The software can export all results to publication-ready quality figures and also allows extraction of data for further user-specific downstream analysis if desired.

The freely available open source code and the underlying mathematical operations that are based on many commonly used packages (e.g. NumPy, SciPy, Matplotlib) will allow expert users to adjust features pipelining the analysis depending on their needs (see Code Availability). The DNN model is trained using Keras/TensorFlow, one of the most popular frameworks for deep learning. While the DNN is pre-trained with DeepFRET, we also provide the option for simulating new data with additional parameters offering the possibility of DNN model re-training to meet the specialized needs of trained users (e.g. multicolour FRET). The programming interface on the other hand allows the convenient implementation of additional scripts pipelining the analysis and to potentially expand it to additional single molecule time series analysis.

## Discussion and Conclusions

smFRET is a powerful toolkit, key for exploring dynamic structural biology, but to meet its full potential, automated standardized and user independent analysis of data are essential. Because the experimental conditions, instruments and biological systems drastically vary across laboratories, the treatment of data based on semi-automatic methods and simplified assumptions could yield different conclusions. DeepFRET is designed to fill this void and analyse data independently of any assumptions and reproducibly across laboratories. Our experiments show that a neural network classifier trained on purely simulated single molecule FRET time series accurately and efficiently recognizes and classifies single molecule FRET both in simulated ground truth data and in real-world dataset. DeepFRET classification accuracy consistently outperformed trace selection using commonly published thresholds. Similarly it supersedes the selection accuracy of human operators and importantly, only requiring a fraction of the time (minutes vs weeks if traces are manually selected). Such drastic reduction of analysis times will allow acquisition of even larger data sets expanding the field for high throughput analysis and improving the robustness of conclusion. The fact that DeepFRET does so only requiring a score threshold, as a sole human intervention, demonstrates its strength as a novel smFRET analysis method and its potential to form a reference against which the quality of the data and the structural biology insights are benchmarked. DeepFRET was found to operate flawlessly for both ALEX and non-ALEX smFRET data highlighting its precise classification and applicability across laboratories and methods. The limited effect of human operators on data selection on the other, hand illustrates its potential to contribute to the standardisation of the field, increasing reproducibility across laboratories. We anticipate that DeepFRET, combined with the advent of commercial single molecule instruments, will contribute in materialising the smFRET as a robust mainstream toolkit for structural biology labs.

DeepFRET’s neural network is trained to operate for smFRET data but our approach of time series classification and sequence annotation can conveniently be extended to consider a spectrum of stochastic single molecule trajectories of individual turnovers including tracking (50) (51–53),constant force measurements (54) and blinking of individual molecules (55, 56) using either simulated or high-quality annotated experimental data for training. Consequently we expect the neural network of DeepFRET or similar approaches to be a paradigm shift and enable new fully automated analysis methodologies related to biomolecular recognition, to protein folding and dynamics and to super resolution. Such advances are paramount for increasing the breadth and impact of single molecule studies to be fully exploited in structural biology.

## Acknowledgements

This work was supported by the Carlsberg Foundation Distinguished Associate Professor program CF16-0797, the VILLUM Foundation Young Investigator Program (grant 10099) and the Villum Foundation Center of Excellence BIONEC (grant 18333) to N.S.H. The Novo Nordisk Foundation Center for Protein Research is supported financially by the Novo Nordisk Foundation (grant NNF14CC0001). This work was also supported by the cryoEM (grant NNF0024386), cryoNET (grant NNF17SA0030214), and Distinguished Investigator (NNF18OC0055061) grants to G.M.G.M. and N.S.H are members of the Integrative Structural Biology Cluster (ISBUC) at the University of Copenhagen.

## Author contributions

J.T performed the method development, implemented the software and evaluated the method with help from M.B.S., T.P., S.B.J., and N.S.H.. S.B.J., B.P. and S.S. helped J.T. to obtain experimental smFRET data. J.T, T.P and N.S.H wrote the manuscript with inputs from G.M., M.B.S. and M.G.M.. N.S.H had the overall strategy and was responsible for the overall project supervision. All authors discussed all data.

## Additional information

correspondence should be addressed to Nikos S. Hatzakis

## Materials and Methods

We first define a nomenclature that will be used throughout the text and plots: *DD*, *DA*, *AA* (donor excitation→donor emission; donor excitation→acceptor emission; acceptor excitation→acceptor emission, respectively). A separate background signal is not considered, as we assume all model inputs to be background-corrected (i.e. background is 0).

### Synthetic smFRET data generation

Deep learning requires large amounts of diverse data in order to generalize well to unseen data. We have developed a method to generate the required thousands of ground-truth traces to cover every type of empirically observed trace, with a dedicated user interface option (Fig. S2). This algorithm includes the generation of TIRF-microscopy smFRET traces of ALEX or non ALEX data. The traces sample any given FRET value with tunable dye photobleaching lifetime, signal noise, dye blinking, donor bleedthrough, aggregates (i.e. more than one donor/acceptor fluorophore) of any given size, as well as well as a “scrambling” feature, to account for fluorophore phenomena that could not be classified as stemming from smFRET.

In order to generate traces, for each pair we first generate the underlying FRET states from an adjustable Hidden Markov model, and assume unscaled unit-intensities for *DD, DA, AA*. Then, if the energy transferred is defined by

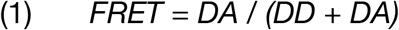

the remaining intensity of the donor is

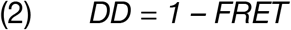

and from (1), the transferred intensity is

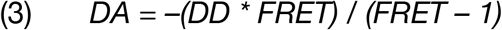

In a perfectly-aligned setup, one can expect that

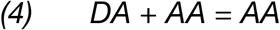

such that the stoichiometry S will be exactly

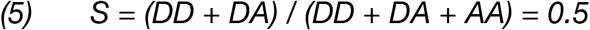

Initially, all fluorophores are simulated with intensity of 1 (with absolute scaling only adjusted after applying all other parameters). Additionally, the intensity of AA should always be 1, regardless of the current FRET state. In practice, the AA intensity may not be exactly half of DD+DA (and consequently one might observe S that deviates slightly from 0.5). To account for this, we uniformly sample “AA-mismatch” as a percentage of the unit intensity signal. Upon fluorophore photobleaching, with lifetimes sampled from an exponential distribution, either DD or DA/AA is set to 0. Noise, AA-mismatch and donor bleedthrough are added to the ground truth signals to obtain the *observable* DD, DA, and AA, we can calculate realistic, *observable* values for E and S. For each synthetic trace, the noise is drawn from a Normal (μ = 0, σ) distribution of varying σ. We found that, on top of the normally distributed noise, we could add the noise from a (centered) Gamma (k = 1, θ = 1.1) distribution multiplied with the noise amplitude at each frame (and is thus controlled via the noise parameter). This did not visually alter the spread of the distribution significantly, but improved robustness of predictions on real data, as we found empirically that the noise of experimental data never exactly followed a pure normal distribution.

State-of-the-art neural networks are able to achieve human-like or better performance on a wide range of classification tasks. Recently, however, it has been demonstrated how small modifications to the input can lead to wildly inaccurate outputs (57). During the development of our smFRET classification model, we observed how photophysical artifacting (described as ‘interesting effects’ by TJ Ha’s group (8)) would lead the model to make confident yet very inaccurate predictions To fix this, our trace generation algorithm contains extensive “scrambling”; we found that by randomly flipping one of the channels, creating strong correlations by multiplication of the channels or adding bursts of high noise and long dark states we could avoid “adversarial-like” predictions. We note that scrambled data is not meant to mimic observable data, but instead to make the model robust against mis-predictions on highly aberrant data that does not fall into the other observable categories.

We generated ground truth traces, where every frame of the sequence was labelled as one of 5 categories: “(B) bleached”, “(A) aggregate”, “(N) noisy”, “(X) scrambled”, “(S) static smFRET” or “(D) dynamic smFRET”(see Fig. S4 for examples). Additionally, we applied label smoothing with a strength of 0.05, as this has been shown to greatly improve model robustness and prediction confidence (58).

For training the model, we set the following parameters (easily adjustable in the interface See Fig. S6):

– Acceptor-only mismatch between 70 % and 130 % of the donor intensity.
– Up to 4 distinct FRET states with a minimum distance of 0.1 FRET between states, with below 0 and 0.2 probability of transitioning from one state to another, at any frame, as determined by a Markov transition matrix.
– 0.15 probability that the trace is an aggregate.
– 0.20 probability that a non-aggregate trace contains photoblinking.
– 0.15 probability that a trace is scrambled, and in this case 0.90 probability that the scrambling is due to incorrectly colocalized fluorophores.
– Donor-bleedthrough between 0 % and 15 % the donor intensity into the acceptor channel.
– Noise drawn from a Normal(0, σ) distribution with σ between 0.01 and 0.30.
– 0.8 probability that the noise has an additional layer of gamma noise on top, to mimic shot noise.
– Individual trace duration of 300 frames.
– Exponentially decaying photobleaching lifetime centered around 500 frames (which will generate a fraction of traces that don’t contain any photobleaching).
– 0.1 probability that the molecule will fall off the surface at a time given by an exponentially decaying lifetime centered around 500 frames (so it might not happen during the time of observation)

With these parameters, directly applicable as input for the algorithm (see Code Availability), we randomly initially generated 250.000 traces of 300 frames each of randomized configurations. We then under-sampled data to balance the labels (as neural network classifiers perform worse if trained on highly class-imbalanced data sets) based on the first frame of each trace. This resulted in approximately 150.000 traces, roughly equally distributed over the 5 possible classes (bleaching being present in most traces naturally ends up accounting for a higher fraction of the overall frames). We used 80% for training the classifier and the remaining 20% for validation. After the training procedure, we generated an additional test set with 33.000 new traces and under-sampled it as previously, to roughly 20.000 traces, and based our statistical analysis on those alone.

We supplied only the raw features DD, DA and AA to the model (or only DD and DA for the non-ALEX-enabled model), where for each trace, signals were normalized to the max of all signals in that trace, so as to preserve the relative intensities between donor and acceptor. In this way, predictions done on individual smFRET traces are fully independent from every other in a given experiment, and also independent from non-standardized instrument intensity units (i.e. “arbitrary units”).

### Neural network model setup and hyperparameters

An LSTM-RNN (long short-term memory recurrent neural network) classifier was implemented in Keras with TensorFlow as backend. The structure of the network (Fig. S1) was inspired by a recent sequence classifier for ECG time series (41) that employs both skip connections and batch normalization as means to prevent overfitting. Here, we replaced the global pooling layer with stride-1 max pooling layers, and added a bidirectional LSTM layer before the final fully connected layer, which we found lead to more temporally causal and context-sensitive predictions (e.g. if the model spots multiple bleaching steps in the beginning of a trace, this information is propagated throughout, so the whole trace can be confidently marked as aggregated).

Each residual block (“Res” in Fig. S1) contains n = 2^x^ filters, where x is 5 and is incremented by 1 at every 4th block. The kernel size k starts from 16 and is reduced by 4 at every 4th Res block, so as to learn larger-scale features and gradually smaller ones. The initial convolution has the same hyper-parameters as the first residual block. A 1 x k convolution is added in each residual block for efficiency (59). To avoid problems with vanishing gradients throughout such a deep model, each residual block keeps a copy of the input vector and adds it to the output vector (denoted by the “+” symbol). The long short-term memory (LSTM) cell is bidirectional and contains 16 units, and has a dropout rate of 0.4 applied to the outputs. For each frame, the outputs are distributed among six different classes by a dense layer with softmax activation.

The model loss was minimized in batches of 32 samples with the Adam optimizer, using the default parameters and the default learning rate of 0.001 The learning rate was decreased by a factor of 10 if validation loss showed no improvement over 2 consecutive epochs. The training was stopped early if no improvement in the validation loss was observed over 5 consecutive epochs. Convolutional kernels were initialized as proposed by (60). Other layer configurations were left at their Keras defaults. The final model output is passed through a softmax layer, thus that for each frame the probabilities between all classes sum up to exactly 1. Further experimentation with optimizers and learning rates showed no significant improvement over the above configuration.

#### Bleaching detection

In order to avoid having single-frame bleaching triggering the remainder of the trace being marked as bleached, we employ a sliding window over the whole trace. In each window, at least 4 out of 7 frames must be marked as bleached with >0.5 probability by the model. If this condition is satisfied, all frames in the window and every frame onwards is marked as bleached, and excluded from the calculation of smFRET confidence. The model predicts with >99% accuracy bleaching (Fig. 3). Additionally, if bleaching happens faster than the first 15 frames, the whole trace is classified as bleached, regardless of model classification, as the DeepFRET score would otherwise end up being artificially inflated (see below).

For stoichiometry-based thresholding (Fig. 2), we employed a similar sliding window, but instead marked frames as bleached if the stoichiometry was outside of the range (0.3, 0.7).

#### Precision and recall

We use precision and recall to quantify classifier performance. These are defined as

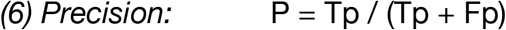

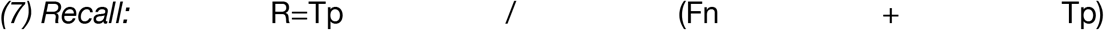

Where Tp, Fp, Fn are True positive, False positive and False negative respectively

#### DeepFRET score calculation and trace classification

In order to calculate the confidence score, probabilities for all categories for each frame are first predicted by the model, and bleached frames (see above) excluded from the score calculation. The average probability p*i* over all frames *t*, for each of the remaining five categories is calculated, resulting in five category scores P*i* for each category *i*.

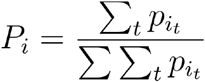

Static smFRET (S), dynamic smFRET (D) scores are summed into the final DeepFRET score, and aggregate (A), noisy (N), and scrambled (X) scores ignored for calculation of this (but retained and displayed for explainability for the user). See Fig. S5, S7 for examples on all trace types.

### Model performance evaluation

#### Noise level of synthetic data

We changed the label of traces to “noisy” if the initial noise was drawn from a Normal(μ = 0, σ) with σ above 0.25. Traces above this level of noise could no longer statistically be approximated as normally distributed by D’Agostino-Pearson two-sided test for normality (Fig. S4) (which is a requirement for fitting the correct number of FRET states in a trace, using a mixture model). Although a σ of 0.20 also fulfilled the p < 0.05 test statistic, we chose to opt for a limit of 0.25, as we found that the neural network would otherwise have a tendency to discard less noisy data too frequently.

#### Trends in human vs. machine selection

To test for differences in the way a human vs. our trained model would select traces, three participants partook in manual selection of generated data (Fig. S15), similar to that of Fig. 2, only this time with 1000 traces, wherein 46 were true smFRET traces and 954 non-usable traces. The number of true smFRET traces and underlying distributions were unknown to the participants.

#### Performance test and comparison with existing software

For testing simple thresholding vs. DeepFRET (Fig. 2, Fig. S12, Fig. S13) we generated data with the following parameters:

– Acceptor-only mismatch between 70 % and 130 % of the donor intensity.
– Donor-bleedthrough of 5 % of the donor intensity into the acceptor channel.
– Noise drawn from a Normal(0.11) distribution
– 1 (0.5 FRET), 2 (0.3, 0.7 FRET) or 3 FRET states (0.2, 0.5, 0.8 FRET)
– Transition probability of 0.1 between states, at each frame.

Other parameters were set to the same value as what is used to generate training data. Furthermore, all generated ground truth traces that didn’t bleach were discarded.

Our definition of “simple thresholding” is based on single molecule intensity, median stoichiometry and the presence of bleaching. We chose here not to use any values for anti-correlation as this assumes that all molecules of interest are equally dynamic, when smFRET studies have shown that this may not always be the case (12, 47, 48).

### Extra features of the software platform

#### Hidden Markov model and statistical analysis

The DeepFRET GUI has the option to fit traces with a Hidden Markov model, with adjustable strictness on the number of states, according to recent best practices for smFRET data analysis, including the ability to switch between predicting states from raw fluorescence intensities or EFRET values. (61) We fit the traces using the pomegranate implementation of the Baum-Welch algorithm (62). We further provide the option to predict state values directly from the Markov Model or from the median of the classified frames for each trace, to maintain compatibility and comparability with current results in the field. We provide clustering of subsequent transition density plots, lifetime estimates with detection of degenerate states, and publication-ready plots for pearson correlation coefficients, DD/DA histograms and EFRET-Stoichiometry histograms.

The Hidden Markov model was verified on externally available data from the kinSoft challenge, as well as simulated data produced within DeepFRET.

#### Data availability

All data used for model training and instructions on how to use it, is available at https://github.com/hatzakislab/DeepFRET-Model/

#### Code availability

We provide DeepFRET as an accessible GUI for everyone, as well as the Python source code for expert users. The code for the GUI as well as compiled executables, with instructions for how to edit and recompile the GUI is located at https://github.com/hatzakislab/DeepFRET-GUI.

## Bibliography

1. E. Lerner, et al., Toward dynamic structural biology: Two decades of single-molecule Förster resonance energy transfer. Science 359, (2018) doi:10.1126/science.aan1133.

2. B. Schuler, W.A. Eaton, Protein folding studied by single-molecule FRET. Curr. Opin. Struct. Biol. 18, 16–26 (2008).

3. B. Hellenkamp, et al., Precision and accuracy of single-molecule FRET measurements-a multi-laboratory benchmark study. Nat. Methods 15, 669–676 (2018).

4. M.F. Juette, et al., Single-molecule imaging of non-equilibrium molecular ensembles on the millisecond timescale. Nat. Methods 13, 341–344 (2016).

5. S. Preus, S.L. Noer, L.L. Hildebrandt, D. Gudnason, V. Birkedal, iSMS: single-molecule FRET microscopy software. Nat. Methods 12, 593–594 (2015).

6. M. Dimura, et al., Quantitative FRET studies and integrative modeling unravel the structure and dynamics of biomolecular systems. Curr. Opin. Struct. Biol. 40, 163–185 (2016).

7. E.D. Holmstrom, Z. Liu, D. Nettels, R.B. Best, B. Schuler, Disordered RNA chaperones can enhance nucleic acid folding via local charge screening. Nat. Commun. 10, 2453 (2019).

8. R. Roy, S. Hohng, T. Ha, A practical guide to single-molecule FRET. Nat. Methods 5, 507–516 (2008).

9. M.D. Newton, et al., DNA stretching induces Cas9 off-target activity. Nat. Struct. Mol. Biol. 26, 185–192 (2019).

10. S. Stella, et al., Conformational Activation Promotes CRISPR-Cas12a Catalysis and Resetting of the Endonuclease Activity. Cell 175, 1856–1871.e21 (2018).

11. S. Kalinin, et al., A toolkit and benchmark study for FRET-restrained high-precision structural modeling. Nat. Methods 9, 1218–1225 (2012).

12. S. Kilic, et al., Single-molecule FRET reveals multiscale chromatin dynamics modulated by HP1α. Nat. Commun. 9, 235 (2018).

13. C. Ratzke, B. Hellenkamp, T. Hugel, Four-colour FRET reveals directionality in the Hsp90 multicomponent machinery. Nat. Commun. 5, 4192 (2014).

14. D. Dulin, et al., Pausing controls branching between productive and non-productive pathways during initial transcription in bacteria. Nat. Commun. 9, 1478 (2018).

15. T.D. Craggs, A.N. Kapanidis, Six steps closer to FRET-driven structural biology. Nat. Methods 9, 1157–1158 (2012).

16. P. Schluesche, G. Stelzer, E. Piaia, D.C. Lamb, M. Meisterernst, NC2 mobilizes TBP on core promoter TATA boxes. Nat. Struct. Mol. Biol. 14, 1196–1201 (2007).

17. S. Sharma, et al., Monitoring protein conformation along the pathway of chaperonin-assisted folding. Cell 133, 142–153 (2008).

18. I.H. Stein, C. Steinhauer, P. Tinnefeld, Single-molecule four-color FRET visualizes energy-transfer paths on DNA origami. J. Am. Chem. Soc. 133, 4193–4195 (2011).

19. J.J. Sakon, K.R. Weninger, Detecting the conformation of individual proteins in live cells. Nat. Methods 7, 203–205 (2010).

20. K. Okamoto, K. Hibino, Y. Sako, In-cell single-molecule FRET measurements reveal three conformational state changes in RAF protein. Biochim. Biophys. Acta Gen. Subj. doi:10.1016/j.bbagen.2019.04.022 (2019) doi:10.1016/j.bbagen.2019.04.022.

21. J. Hon, R.L. Gonzalez, Bayesian-Estimated Hierarchical HMMs Enable Robust Analysis of Single-Molecule Kinetic Heterogeneity. Biophys. J. 116, 1790–1802 (2019).

22. M. Greenfeld, D.S. Pavlichin, H. Mabuchi, D. Herschlag, Single Molecule Analysis Research Tool (SMART): an integrated approach for analyzing single molecule data. PLoS ONE 7, e30024 (2012).

23. S. Schmid, M. Götz, T. Hugel, Single-Molecule Analysis beyond Dwell Times: Demonstration and Assessment in and out of Equilibrium. Biophys. J. 111, 1375–1384 (2016).

24. W.R. Algar, N. Hildebrandt, S.S. Vogel, I.L. Medintz, FRET as a biomolecular research tool – understanding its potential while avoiding pitfalls. Nat. Methods 16, 815–829 (2019).

25. E. Lerner, et al., The FRET-based structural dynamics challenge -- community contributions to consistent and open science practices. (2020).

26. A. Sali, et al., Outcome of the First wwPDB Hybrid/Integrative Methods Task Force Workshop. Structure 23, 1156–1167 (2015).

27. Y. LeCun, Y. Bengio, G. Hinton, Deep learning. Nature 521, 436–444 (2015).

28. T.A. collaboration, A neural network clustering algorithm for the ATLAS silicon pixel detector. J. Inst. 9, P09009–P09009 (2014).

29. P. Zhang, et al., Analyzing complex single-molecule emission patterns with deep learning. Nat. Methods 15, 913–916 (2018).

30. W. Ouyang, A. Aristov, M. Lelek, X. Hao, C. Zimmer, Deep learning massively accelerates super-resolution localization microscopy. Nat. Biotechnol. 36, 460–468 (2018).

31. E.M. Christiansen, et al., In silico labeling: predicting fluorescent labels in unlabeled images. Cell 173, 792–803.e19 (2018).

32. T. Falk, et al., U-Net: deep learning for cell counting, detection, and morphometry. Nat. Methods 16, 67–70 (2019).

33. D.T. Jones, Setting the standards for machine learning in biology. Nat. Rev. Mol. Cell Biol. 20, 659–660 (2019).

34. S. Berg, et al., ilastik: interactive machine learning for (bio)image analysis. Nat. Methods 16, 1226–1232 (2019).

35. J.T. Smith, et al., Fast fit-free analysis of fluorescence lifetime imaging via deep learning. Proc Natl Acad Sci USA doi:10.1073/pnas.1912707116 (2019) doi:10.1073/pnas.1912707116.

36. P.A. Gómez-García, E.T. Garbacik, J.J. Otterstrom, M.F. Garcia-Parajo, M. Lakadamyali, Excitation-multiplexed multicolor superresolution imaging with fm-STORM and fm-DNA-PAINT. Proc Natl Acad Sci USA 115, 12991–12996 (2018).

37. S.J. Yang, et al., Assessing microscope image focus quality with deep learning. BMC Bioinformatics 19, 77 (2018).

38. A.C.-Y. Wu, S.A. Rifkin, Aro: a machine learning approach to identifying single molecules and estimating classification error in fluorescence microscopy images. BMC Bioinformatics 16, 102 (2015).

39. Y. Matsunaga, Y. Sugita, Linking time-series of single-molecule experiments with molecular dynamics simulations by machine learning. elife 7, (2018) doi:10.7554/eLife.32668.

40. R.P. Thomsen, et al., A large size-selective DNA nanopore with sensing applications. Nat Commun (2019).

41. A.Y. Hannun, et al., Cardiologist-level arrhythmia detection and classification in ambulatory electrocardiograms using a deep neural network. Nat. Med. 25, 65–69 (2019).

42. R.R. Wick, L.M. Judd, K.E. Holt, Deepbinner: Demultiplexing barcoded Oxford Nanopore reads with deep convolutional neural networks. PLoS Comput. Biol. 14, e1006583 (2018).

43. F. Karim, S. Majumdar, H. Darabi, S. Chen, LSTM fully convolutional networks for time series classification. IEEE Access 6, 1662–1669 (2018).

44. S.L. Oh, E.Y.K. Ng, R.S. Tan, U.R. Acharya, Automated diagnosis of arrhythmia using combination of CNN and LSTM techniques with variable length heart beats. Comput. Biol. Med. 102, 278–287 (2018).

45. N.K. Lee, et al., Accurate FRET measurements within single diffusing biomolecules using alternating-laser excitation. Biophys. J. 88, 2939–2953 (2005).

46. J. Hohlbein, T.D. Craggs, T. Cordes, Alternating-laser excitation: single-molecule FRET and beyond. Chem. Soc. Rev. 43, 1156–1171 (2014).

47. S. Osuka, et al., Real-time observation of flexible domain movements in CRISPR-Cas9. EMBO J. 37, (2018) doi:10.15252/embj.201796941.

48. S. He, C. Yang, S. Peng, C. Chen, X.S. Zhao, Single-molecule study on conformational dynamics of M.HhaI. RSC Adv. 9, 14745–14749 (2019).

49. S. Wood, A.R. Ferré-D’Amaré, D. Rueda, Allosteric tertiary interactions preorganize the c-di-GMP riboswitch and accelerate ligand binding. ACS Chem. Biol. 7, 920–927 (2012).

50. S.S.-R. Bohr, et al., Direct observation of Thermomyces lanuginosus lipase diffusional states by Single Particle Tracking and their remodeling by mutations and inhibition. Sci. Rep. 9, 16169 (2019).

51. H.P. Lu, L. Xun, X.S. Xie, Single-molecule enzymatic dynamics. Science 282, 1877–1882 (1998).

52. F. Persson, M. Lindén, C. Unoson, J. Elf, Extracting intracellular diffusive states and transition rates from single-molecule tracking data. Nat. Methods 10, 265–269 (2013).

53. L.S. Ferro, S. Can, M.A. Turner, M.M. ElShenawy, A. Yildiz, Kinesin and dynein use distinct mechanisms to bypass obstacles. elife 8, (2019) doi:10.7554/eLife.48629.

54. D.H. Goldman, et al., Ribosome. Mechanical force releases nascent chain-mediated ribosome arrest in vitro and in vivo. Science 348, 457–460 (2015).

55. N. Durisic, L. Laparra-Cuervo, A. Sandoval-Álvarez, J.S. Borbely, M. Lakadamyali, Single-molecule evaluation of fluorescent protein photoactivation efficiency using an in vivo nanotemplate. Nat. Methods 11, 156–162 (2014).

56. G. Wang, et al., Structural plasticity of actin-spectrin membrane skeleton and functional role of actin and spectrin in axon degeneration. elife 8, (2019) doi:10.7554/eLife.38730.

57. I. Goodfellow, J. Shlens, C. Szegedy, Explaining and Harnessing Adversarial Examples. arXiv 1412.6572v3 [stat.ML] (2014).

58. A. Shafahi, A. Ghiasi, F. Huang, T. Goldstein, Label Smoothing and Logit Squeezing: A Replacement for Adversarial Training? arXiv (2019).

59. K. He, X. Zhang, S. Ren, J. Sun, “Deep residual learning for image recognition.” in IEEE Conference on Computer Vision and Pattern Recognition (CVPR) (IEEE, 2016), pp. 770–778.

60. K. He, X. Zhang, S. Ren, J. Sun, “Delving Deep into Rectifiers: Surpassing Human-Level Performance on ImageNet Classification.” in 2015 IEEE International Conference on Computer Vision (ICCV) (IEEE, 2015), pp. 1026–1034.

61. D. Kelly, M. Dillingham, A. Hudson, K. Wiesner, A new method for inferring hidden markov models from noisy time sequences. PLoS ONE 7, e29703 (2012).

62. J. Schreiber, pomegranate: Fast and Flexible Probabilistic Modeling in Python. Journal of Machine Learning Research 18, 1–6 (2018).

