## Supplementary figures for "DeepFRET: Rapid and automated single molecule FRET data classification using deep learning"

**Figure Supplement**

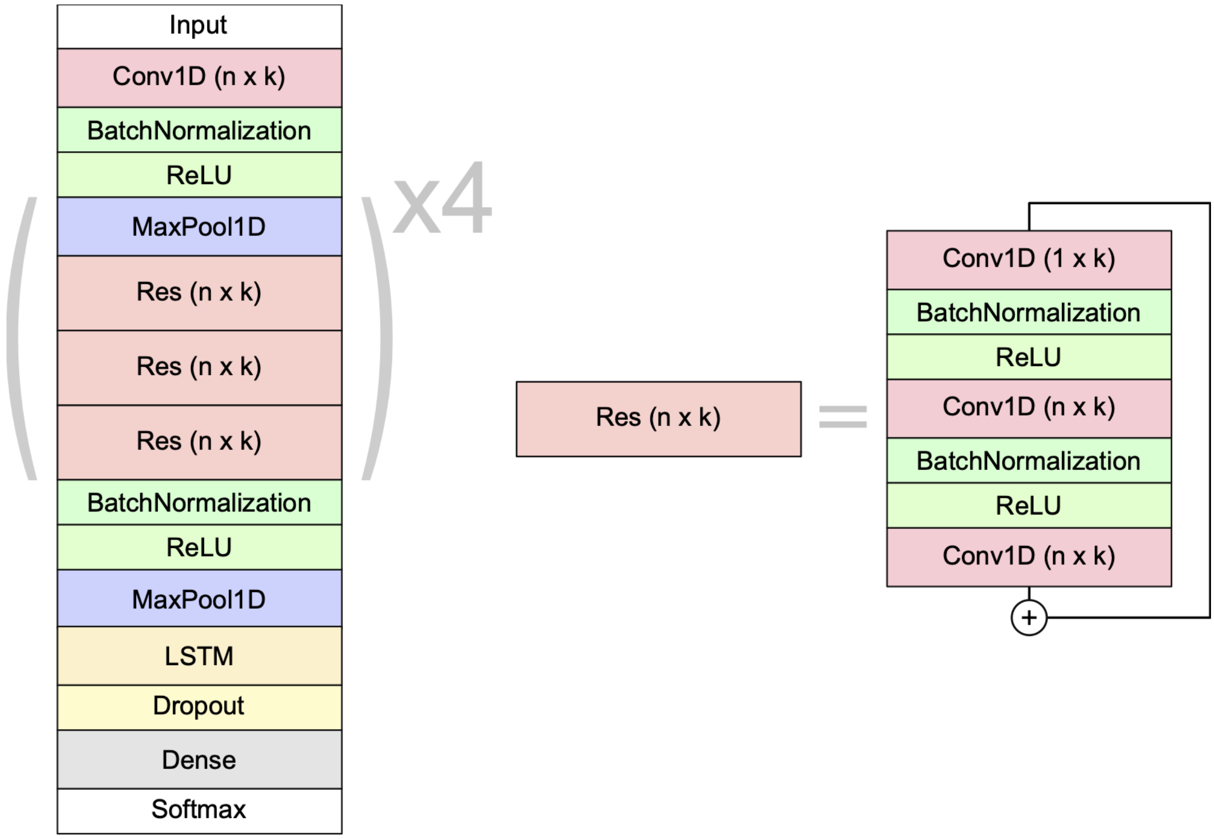


**Figure S1 | Schematic overview of the neural network model architecture.** Each residual block ("Res") is made up of 1D convolutional filters of size n x k with batch normalization and ReLU activations in between. The initial convolution after the input layer has the same hyper-parameters as the first residual block. A 1 x k convolution is added in each bottleneck unit for efficiency. The "+" symbol denotes a skip-connection, where the residual block's input is added to its output. The final convolved output goes to a bidirectional long short-term memory (LSTM) layer. For each frame, the outputs are distributed among the different classes by a dense layer with softmax activation.

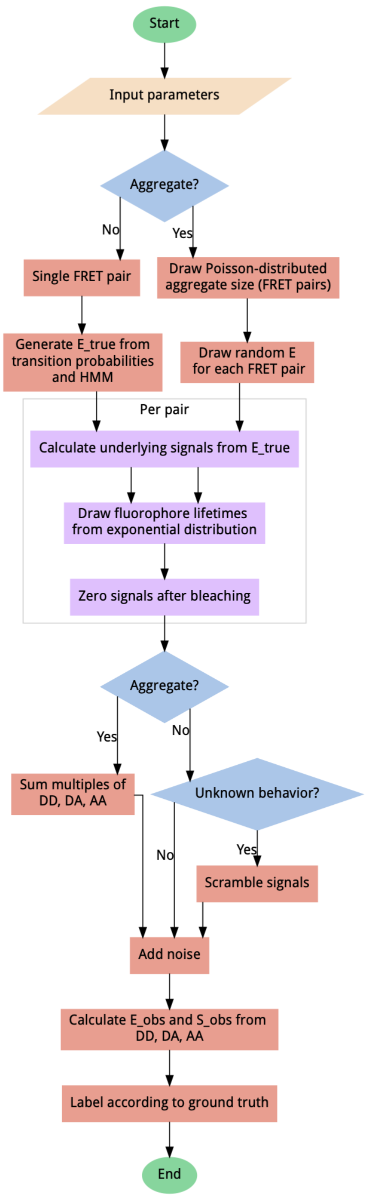


**Figure S2 | Data generator algorithm overview.**

To generate unbiased distributions of smFRET traces, most steps include randomization, sampling from uniform distributions (except for bleaching probability, which follows an exponential distribution) for the given input parameter ranges. User-adjustable parameters include the probability of aggregation, fluorophore blinking, bleaching and more. All output data is automatically sequence-annotated with the ground truth behaviour, for supervised machine learning. A scrambling probability is used to generate a fraction of heavily distorted data to improve model robustness and prevent misclassification of data containing artefacts.

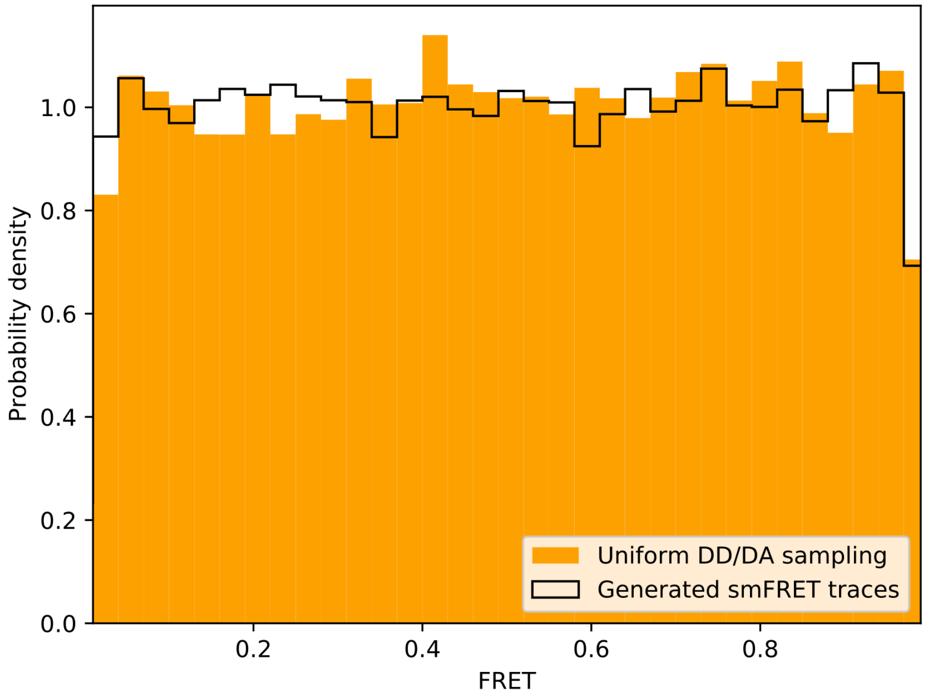


**Figure S3 | FRET values are uniformly sampled for all ground truth smFRET traces.** smFRET traces were generated with either 1, 2, or 3 randomly defined FRET states with the same transition probabilities as used for model training. Comparing FRET calculated from uniformly sampled donor/acceptor intensities between 0 and 1 reveals identical, uniform distributions.  The simulated training data does not introduce any FRET-state bias in the model training.

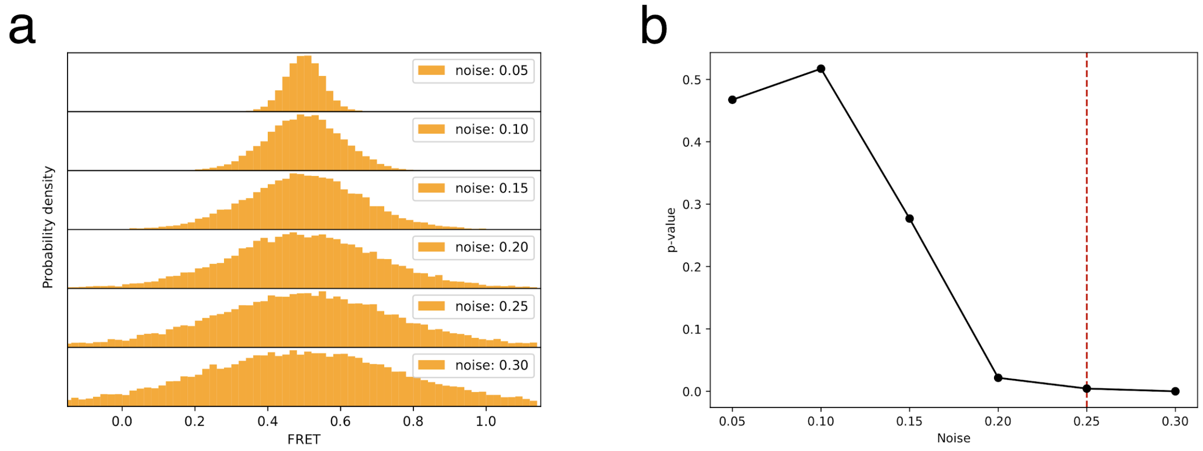


**Figure S4 | Noise threshold for simulated data.** a) To tune the accepted noise level for our deep learning model, we simulated 200 smFRET traces with a single state at 0.5 FRET, with varying noise levels. b) At a noise level of 0.25, the distributions in a) could no longer be considered normal, using a D'Agostino-Pearson test for normality ($p < 0.05$).

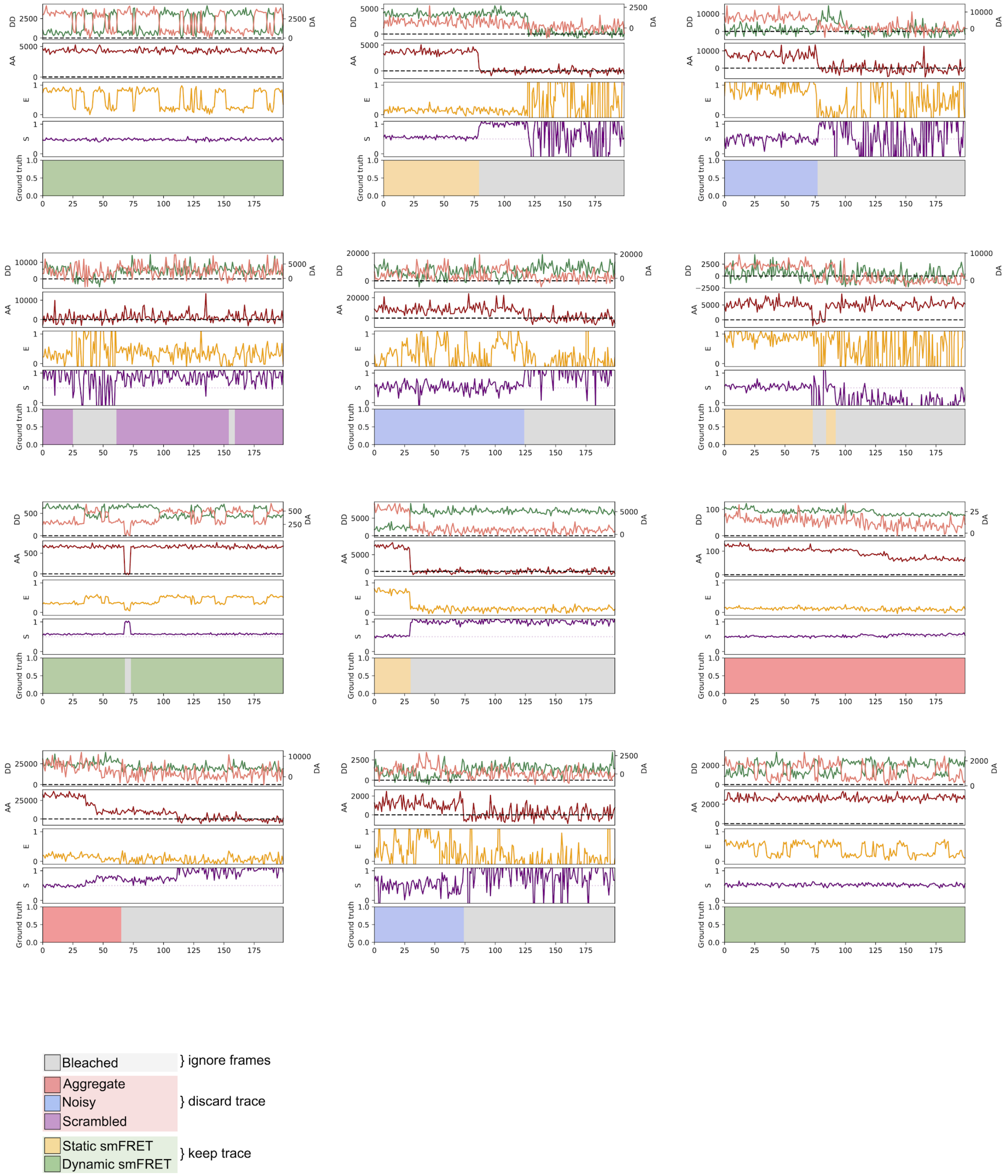


**Figure S5 | Examples of randomly generated traces.**

Randomly generated traces, corresponding to each of the six defined categories, shown with raw signals, E (FRET) and S (stoichiometry). The colour code in the bottom panel corresponds to the frame-level ground truth labeling of each trace.

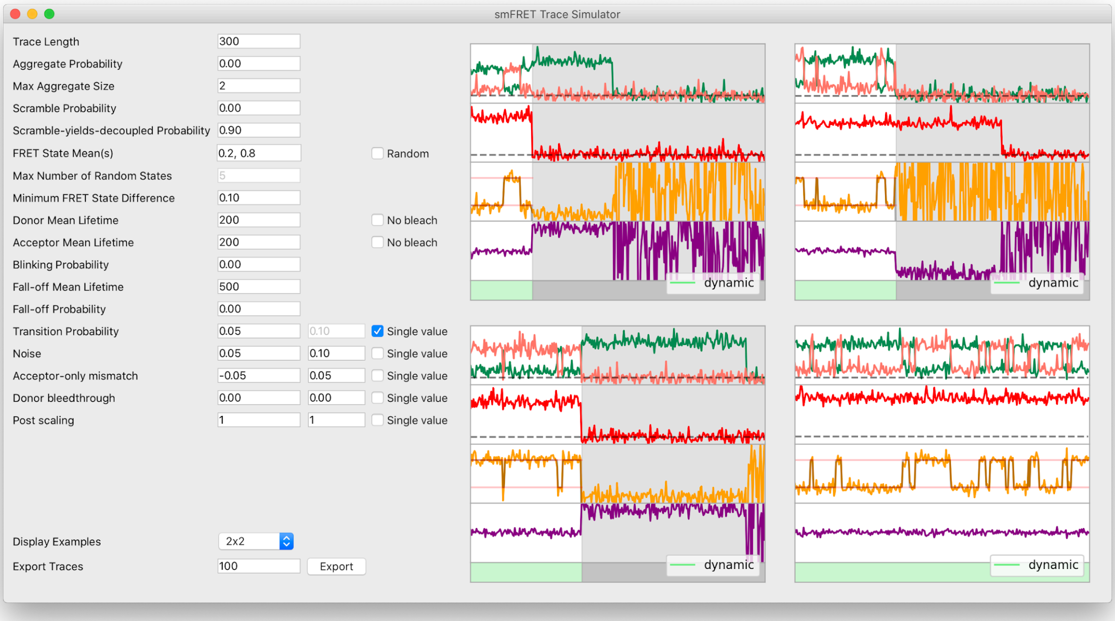


**Figure S6 | Trace simulator interface.** To facilitate fast and easy experimentation of any kind on smFRET traces, the DeepFRET GUI includes a visual trace simulator with ground-truth labels associated with every trace. The traces can be exported to ASCII or DAT and used for all kinds of statistical procedures.

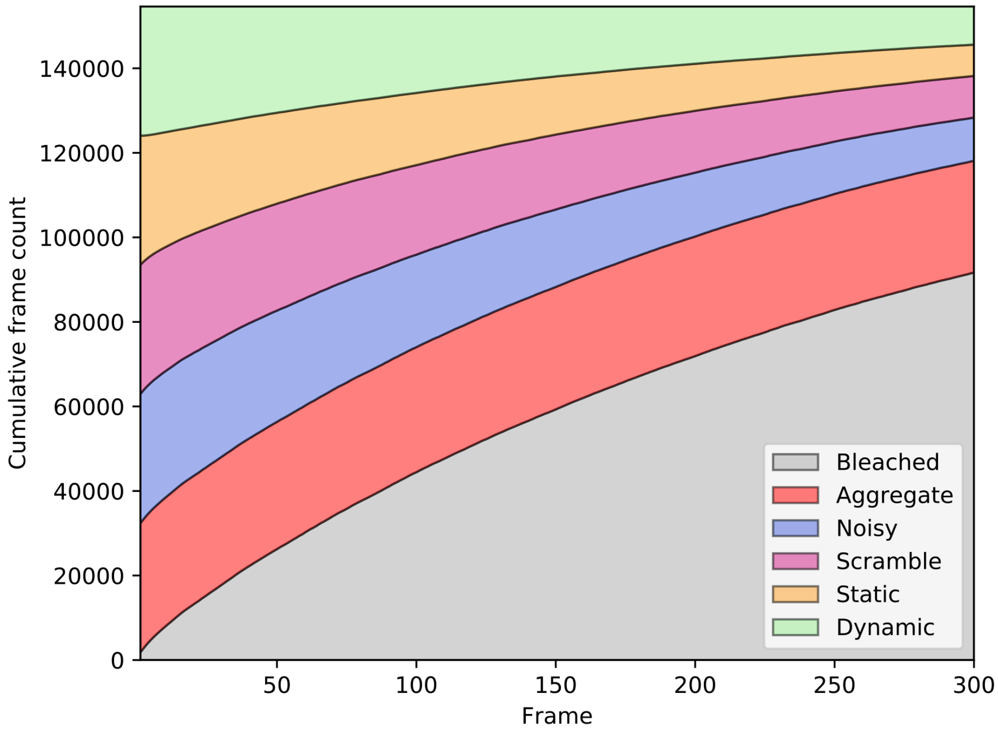


**Figure S7 | Training label dependence on the frame number.**
The distribution of labels for all frames in the training dataset, after balancing classes based on the first frame. The “bleaching” label is excluded from balancing, as it occurs in most of the generated traces, due to photobleaching.

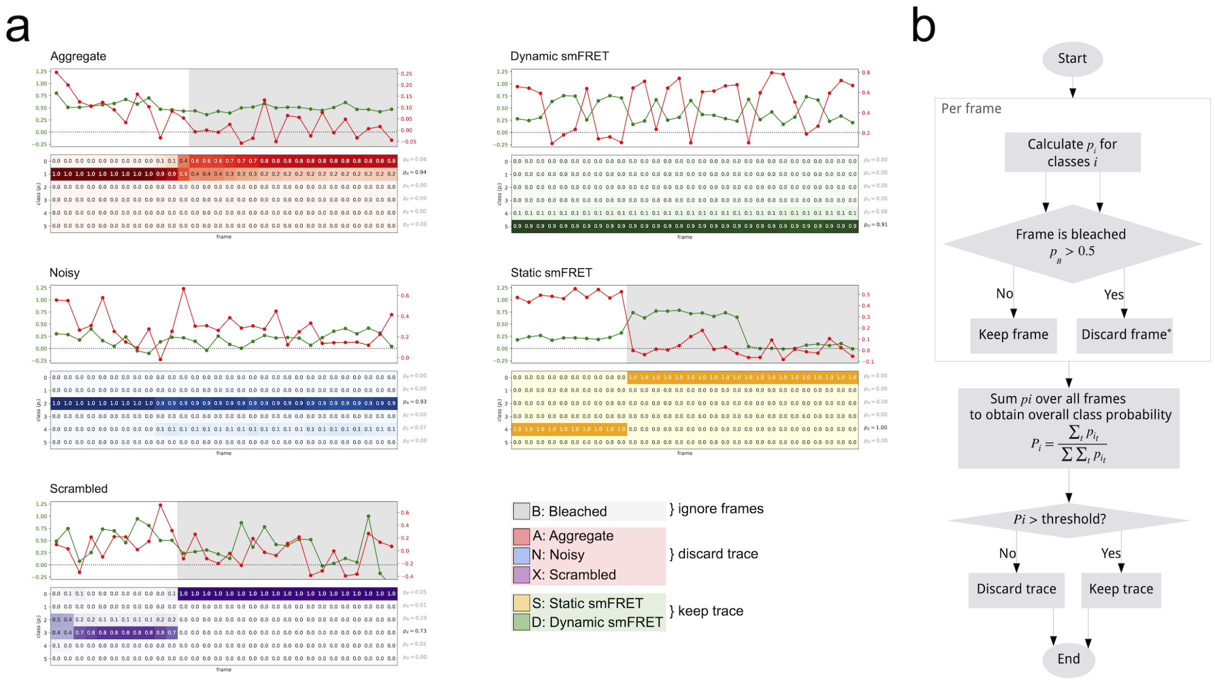


**Figure S8 | Calculation of confidence score from model predictions.** a) Short toy-examples of selected traces for each category and the corresponding frame-wise predictions used for calculating confidence scores. Colour code corresponds to classified behavior. All bleached data points (marked with grey in the trace) are excluded from the confidence score calculation. b) Step-by-step calculation of trace scores.

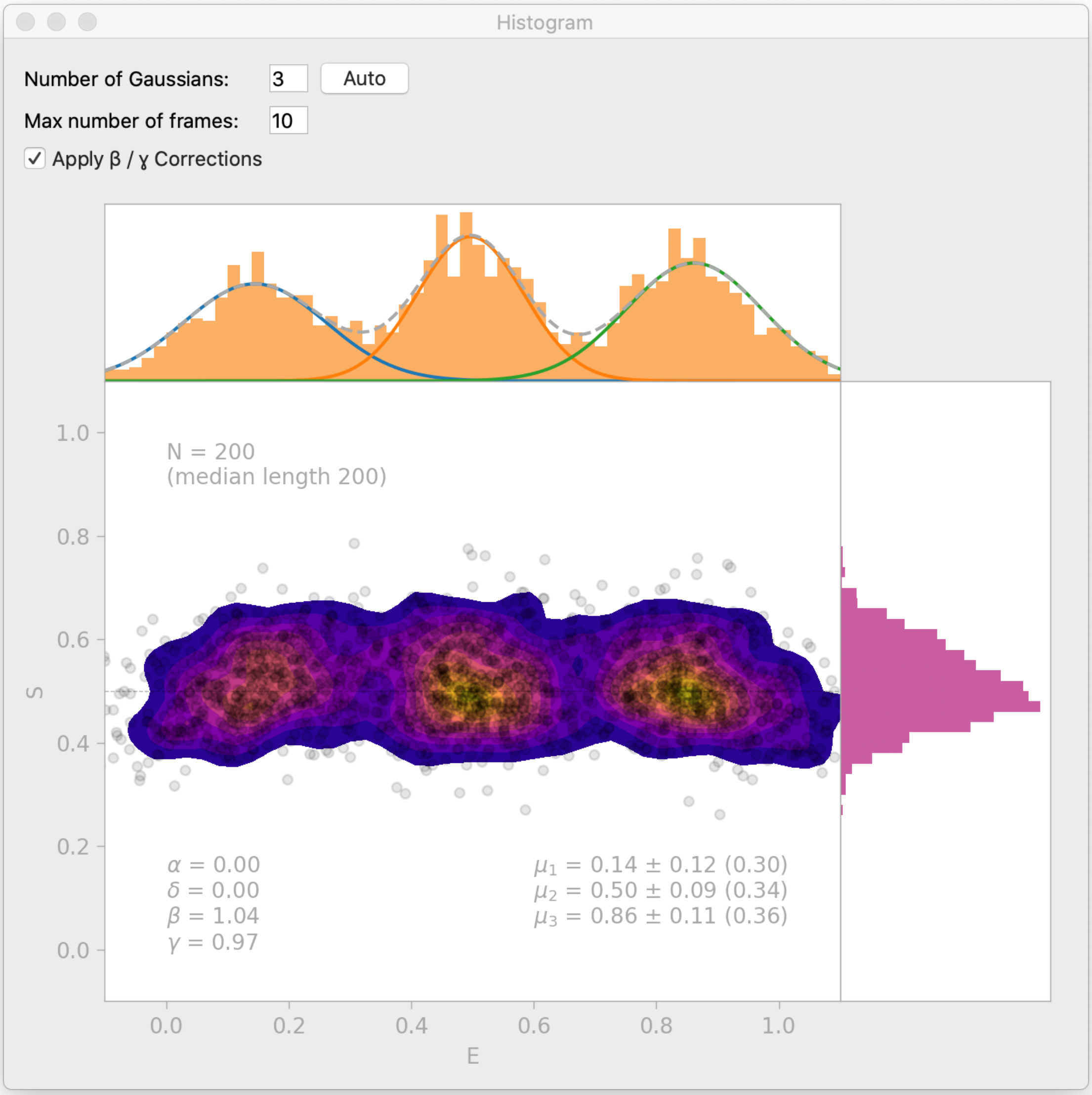


**Figure S9 | Histogram interface window.** DeepFRET includes a histogram window which automatically finds BIC-optimal Gaussian Mixtures, and allows the user to test enter the number of Gaussians to include in a Mixture model. The Histogram Interface also includes an option for β- and γ-correction. Data is simulated from DeepFRETs trace generator with 3 states with means 0.15, 0.5 and 0.85, as in Fig. S13b.

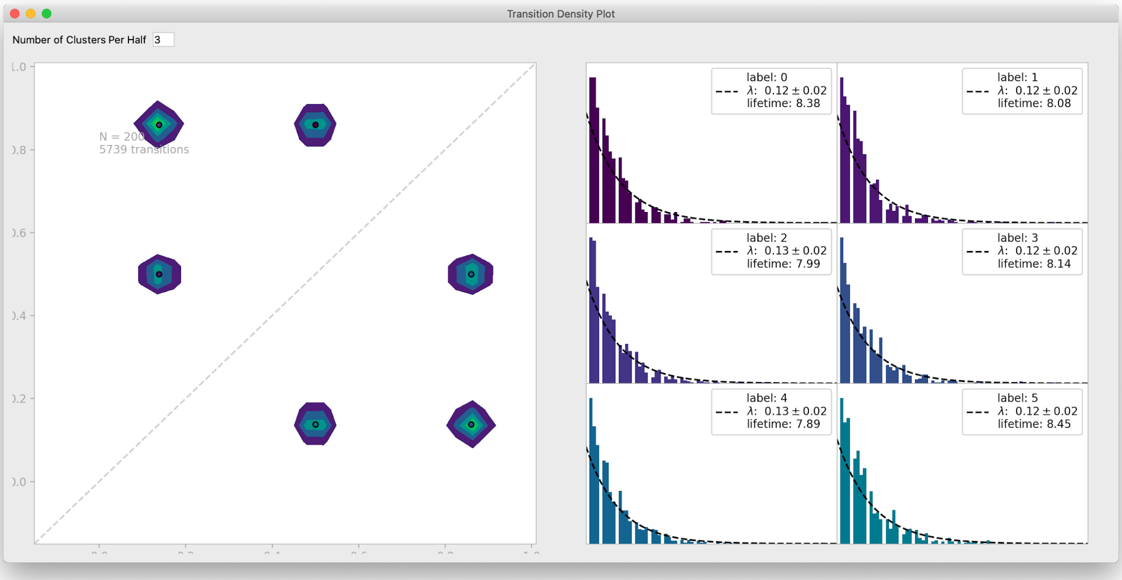


**Figure S10 | Transition Density window.** To visualize transitions between FRET states, DeepFRET includes a window to fit exponential lifetime distributions of clusters in a Transition Density plot. The Transition Density interface also allows the user to test a different number of clusters for the Transition Density Window. Data is simulated from DeepFRETs trace generator with 3 states with means 0.15, 0.5 and 0.85, as in Fig. S13b.

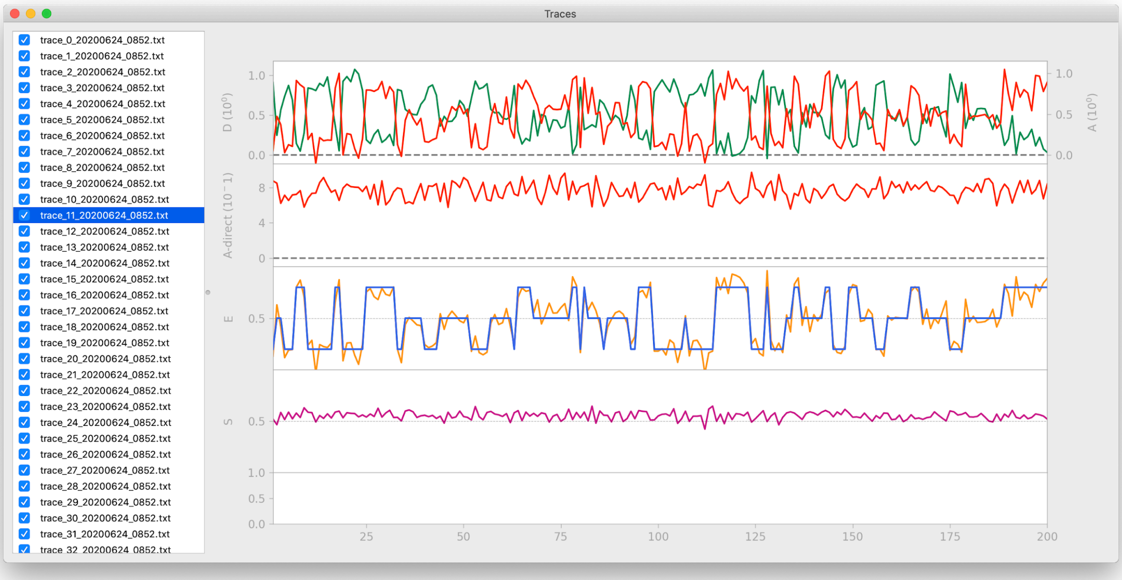


**Figure S11 | Traces Window** To provide a direct visualization of the Hidden Markov Modelling of smFRET traces, traces can be imported directly for HMM analysis using the commonly used pomegranate implementation of the Baum-Welch algorithm. The transitions from the Traces window are used in the Transition Density Window. Data is simulated from DeepFRETs trace generator with 3 states with means 0.15, 0.5 and 0.85, as in Fig. S13b.

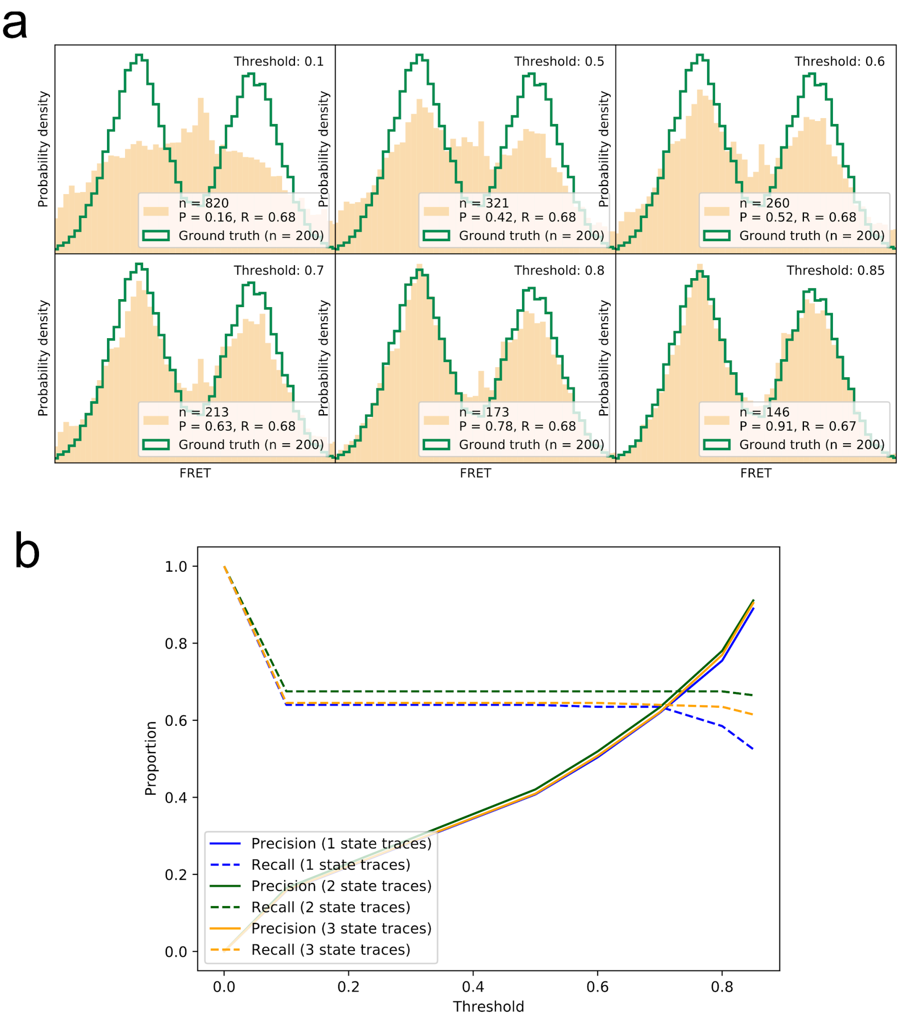


**Figure S12 | Precision-recall for correctly identifying smFRET traces at different thresholds. a)** Visual display of recovered smFRET distributions for several thresholds for a 2-state system (same data as Fig. 2)**. b)** The precision and recall are calculated for all thresholds on the datasets shown in Fig. 2 and Fig. S8. For 1-, 2- and 3-state systems we find that a threshold around 0.7-0.8 provides the best tradeoff between the two measures, in order to recover the underlying FRET distribution faithfully.

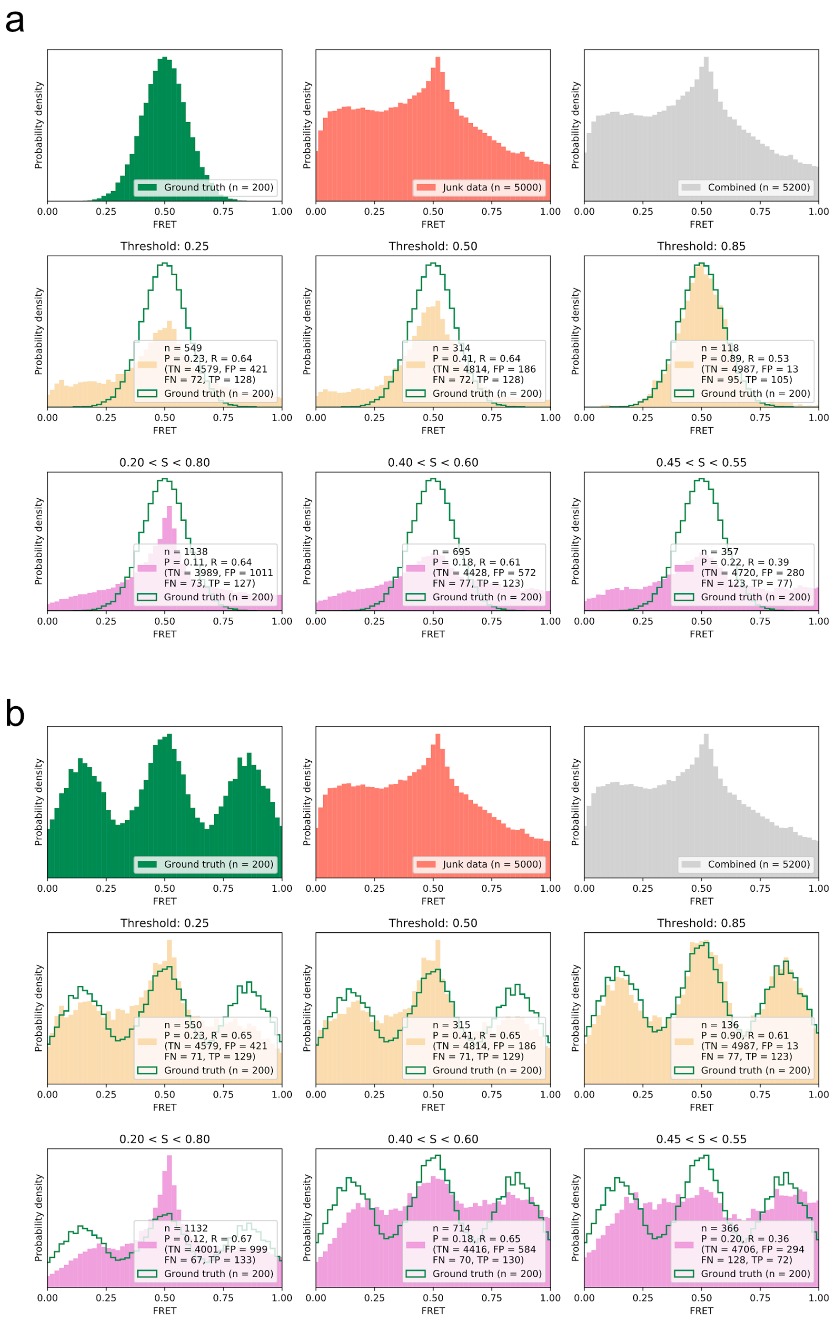


**Figure S13 | Comparison of smFRET distribution recovery by DeepFRET at different thresholds and semi-automated methodologies under various conditions in the absence of further human intervention.** In each of the figures (a) and (b); Top: ground truth distribution and distribution of randomly selected non-smFRET data from an external test set. Middle: Performance of DeepFRET sorting at different thresholds. Bleached frames were excluded from analysis in all cases. Bottom: Performance of sorting by semi-automated methodologies using common thresholds-based sorting at different thresholds. Classification metrics for smFRET trace detection are shown in the graph. **a)** The performance, given a 1-state system centered at FRET mean value 0.5. **b)** The performance, given a 3-state system centered at FRET mean values 0.15, 0.5 and 0.85.

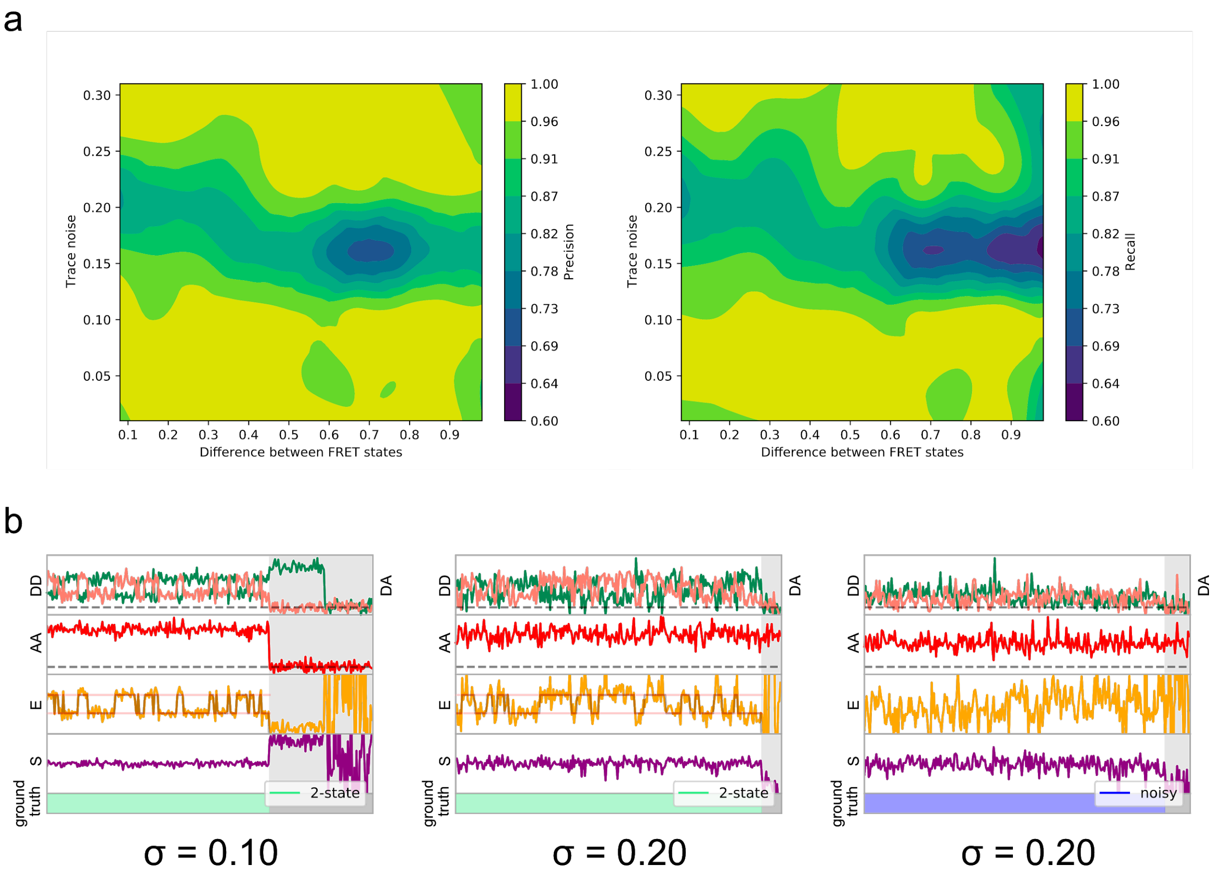


**Figure S14 | Relationship between model performance and noise level.** a) 3D illustration of the Precision or recall (pseudocolour) as a function of trace noise  and the distance between FRET states for dynamic 2-state traces. Prediction accuracy is minimal at ~0.2 noise levels. At lower noise the trace is accepted and correctly classified with an accuracy of >0.96, while at high noise levels the trace is accurately classified as non FRET. . b) Examples of simulated dynamic traces used, with input simulation noise. At a high simulated noise level, only some traces are below the lower limit of acceptable noise, when measuring the per-state standard deviation of FRET of the final output trace.

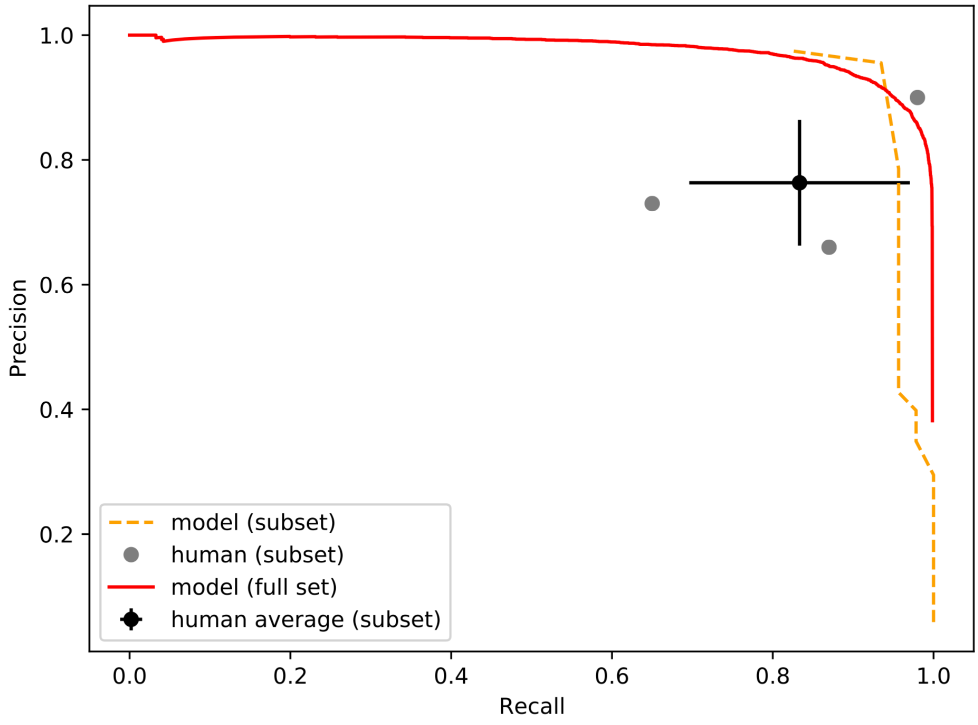


**Figure S15 | Precision-recall of the neural network and human participants.** Precision is plotted against recall for all precision on a simulated dataset containing 46 true smFRET traces and 954 non-smFRET traces. Additionally, the precision-recall for the entire test set (20,000 traces) is plotted for the model. The error bars on the average participant performance represent the standard deviation of the three individual participants.

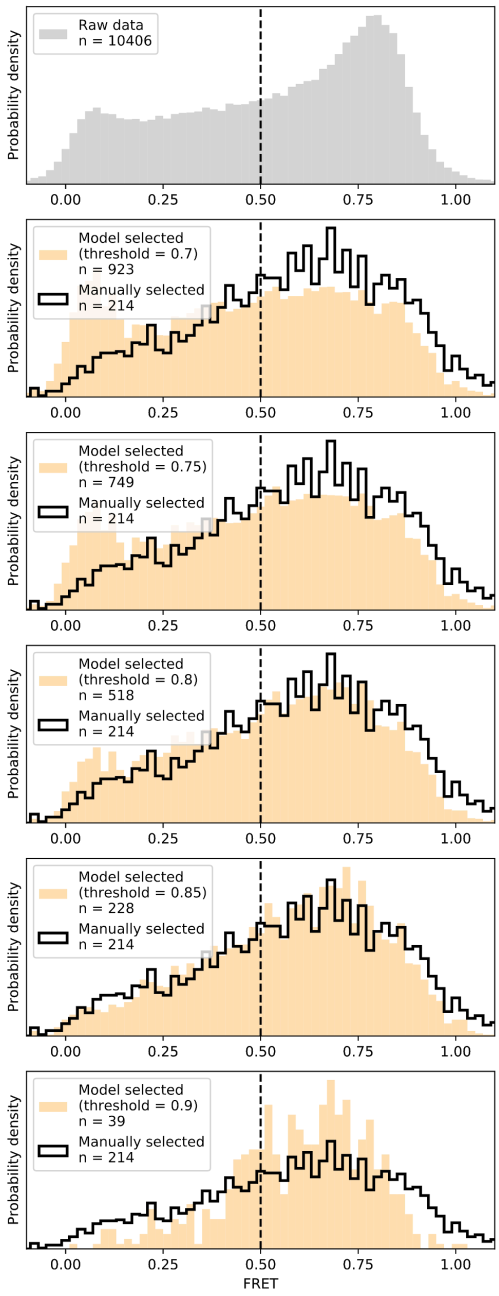


**Figure S16 | DeepFRET applied to experimentally-obtained data.** We applied DeepFRET to a previously obtained smFRET dataset on CRIPSR-Cas12a. At DeepFRET threshold of ~0.8-0.85 an excellent agreement with previously used sorting methodologies involving manual inspection of data was achieved.

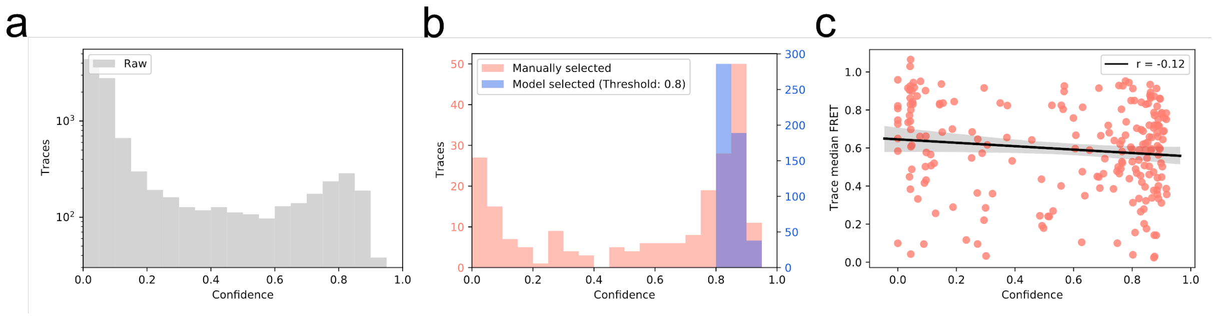


**Figure S17 | Distribution of trace quality of the experimental data set. a)** The predicted confidence score of every trace in experimentally recorded dataset **b)** Confidence score distribution of manual vs. automatic selection of smFRET traces. Automatic selection exclusively accepts high-confidence traces. **c)** Pearson's test displaying no correlation (r = 0.01) between predicted trace confidence and mean FRET in the manually selected dataset.
